# Controlling Heterogeneity and Increasing Titer from Riboswitch-Regulated *Bacillus subtilis* Spores for Time-Delayed Protein Expression Applications

**DOI:** 10.1101/592659

**Authors:** Denis Tamiev, Alyssa Lantz, Grace Vezeau, Howard Salis, Nigel F. Reuel

## Abstract

Sporulated cells have potential as time-delayed expression chassis of proteins for applications such as ‘on-demand’ biologics production, whole cell biosensors, or oral vaccines. However, the desired attributes of high expression rates and low product variances are difficult to maintain from germinated spores. In this work we study the effect of an integrating *vs.* theta replicating plasmid in a wild-type *Bacillus subtilis* and two PolY mutants. The cells were engineered to produce a fluorescent reporter protein (RFP) under the control of a riboswitch activated by theophylline. This allowed for greater sensitivity to point mutations. The fluorescence and cell growth curves were fit with a custom kinetic model and a peak kinetic rate (LKP_max_) was extracted for each clonal population (n = 30 for all cell, vector, and growth combinations). Plasmid based expression yields higher (8.7x) expression rates due to an increased copy number of the expression cassette (10x over integrated). The variance of LKP_max_ values increased 2.07x after sporulation for the wild type strain. This increase in variance from sporulation is very similar to what is observed with UV exposure. This effect can be partially mitigated by the use of PolY knockouts observed in suspended cell growths and adherent biofilms.

---

The ability to express recombinant proteins from engineered cells has revolutionized biotechnology research and biologic therapy production^1^. In most cases, the designed protein product is produced in large batches, purified, and stored for future use. Although much work has gone into the shelf-stabilization of proteins, such as lyophilization, addition of anionic gold particles or storage in polyethylene glycol, these methods will never completely eliminate short shelf-life of purified biological drugs especially when cold-chain storage is not available.^2,3^ An alternative strategy to increasing the shelf-life, is to produce the proteins on-demand, thus circumventing the need to store the product. Certain use cases, such as deep space missions or remote field locations, have spurred interest in time-delayed protein expression solutions^4,5^. One approach is the use of cell extract for *in vitro* protein production; however, these solution-based or lyophilized components suffer from the same shelf-stability concerns as the purified protein, with maximum viability demonstrated at one year^6^. Beyond that, cell free approaches would be difficult to deploy into closed systems, such as soil monitoring, as they require intervention to mix the needed reagents. A better approach for closed systems would be a viable cell that can remain dormant for prolonged periods of time until they are activated with a specific chemical cue to produce a reporter protein signal (color, luminescence, hydrolytic activity, *etc.*). Spores can act in this manner as time-delayed expression chassis.

Sporulation is an important safety mechanism utilized by a phylum of gram-positive bacteria called Firmicutes^7,8^. When resources are limited, they have the ability to adopt a nearly dormant endospore form in which a thick protein coat preserves the genetic information for extended periods of time^9^. Once growth-supporting conditions return, these cells will revive and resume normal cellular processes, including protein expression, metabolism, growth, and self-replication. Successful revival has been observed from truly ancient spores, some formed 25 million years ago^10^. This ability to harbor genetic information for time-delayed protein expression makes endospore-forming cells prime candidates for 1) ‘on-demand,’ time-delayed protein expression, 2) point-of-care (bio)-logic sensors, and 3) oral vaccinations that can germinate *in vivo* (Fig 1A). Some eukaryotic cells, such as yeast, can also form spores and may also be considered candidates for these time-delayed applications; however, the shelf-life of common industrial yeast strains is limited^11^. Of the spore forming bacteria, *B. subtilis* was selected due to its lack of pathogenicity or endotoxins (generally recognized as safe or GRAS organism)^12^ and it is the most common species used for industrial protein expression. Although there are previous studies that have demonstrated the use of *B. subtilis* spores for whole cell biosensors and vaccine delivery ^13,14^, they have all overlooked a critical limitation in spore usage: the heterogeneity of protein expression due low fidelity DNA replication during late stationary growth phase. The heterochronicity of sporulation out growth has been recently highlighted in another study^15^, but a study that assesses the effect of sporulation on expression rates is lacking. Such heterogeneity would impact the proposed time-delay applications, causing variations in sensor calibrations or vaccine dosing. Moreover, it could decrease the production of active protein product, as spores with compromised expression cassette sequences would produce inactive product. Therefore, studying the factors impacting the heterogeneity and titer of protein production from *B. subtilis* spores is the focus of this work.

**Figure 1.**
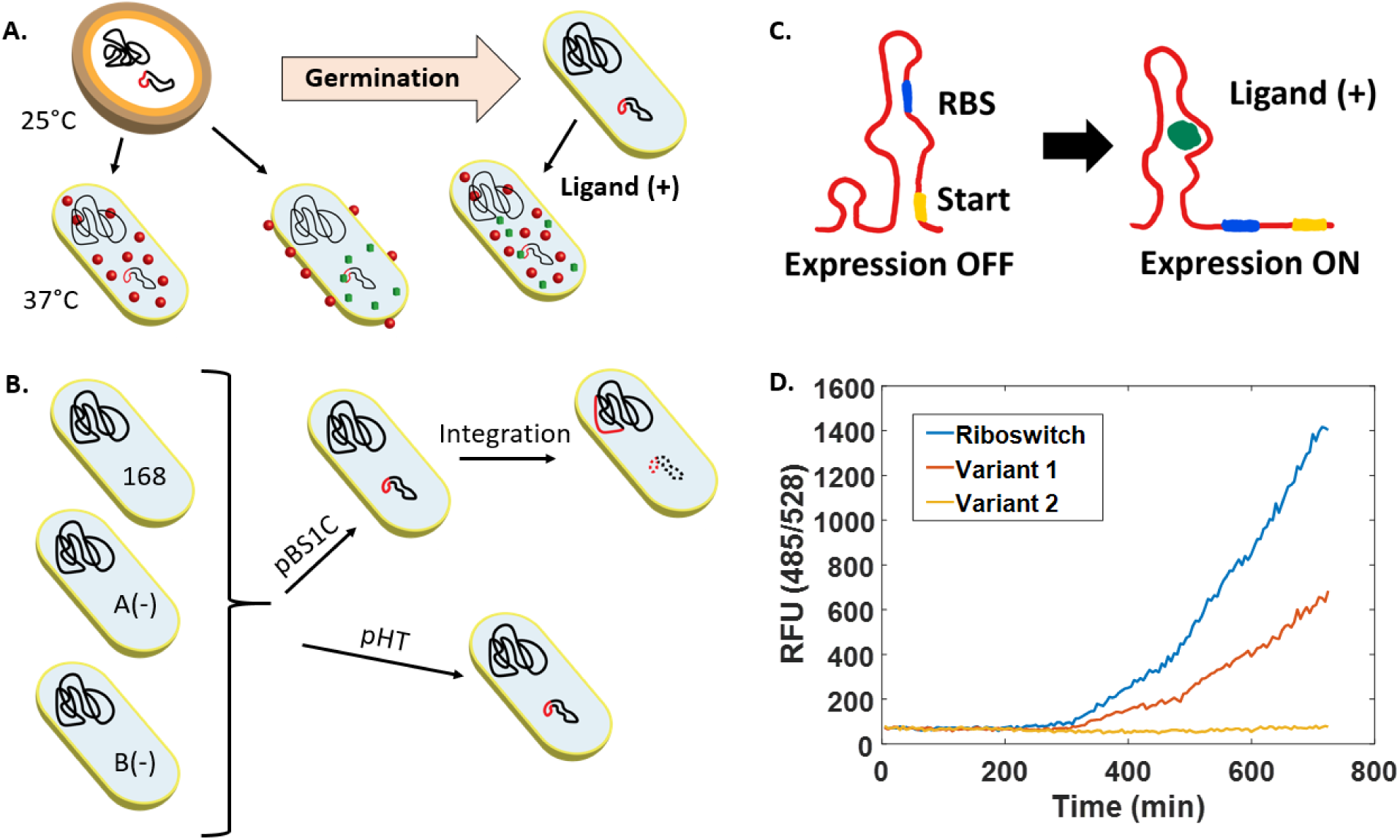
(**A**) Proposed time-delay expression applications for spores: production of biologics, oral vaccines, and whole cell biosensors. (**B**) *Bacillus subtilis* strains used in this study: wild type strain, PolY A(-) and PolY B(-) mutants; also showing genome integrated plasmids (pBS1C) and theta replicating plasmids (pHT) compared in this work. (**C**) Schematic showing regulation of reporter protein (RFP) expression with a riboswitch activated by Theophylline ligand. (**D**) RFP expression from RFP/Theophylline riboswitch and two sequence variants (55.3% and 44.7% sequence identity to the riboswitch, respectively) showing sensitivity to point mutations.

In this study we compare two modes of expression, one directed by a chromosomal integrating plasmid (pBS1C) and a second using a theta replicating plasmid (pHT)^16^ (Fig 1B). Legacy *B. subtilis* protein expression techniques rely on homology-driven, chromosomal-integrating plasmids (episomes), while typically, theta replicating plasmids are reserved for protein expression in *E. coli*. The stated reason is low sequence stability of theta replicating plasmids^16^. We hypothesized that although the integrating plasmid would produce better homogeneity, it would be at the expense of protein titer because of the stoichiometry of the plasmid copy number as well as the accessibility of DNA to low fidelity polymerases. Herein we determine if post-sporulation protein expression with stable, theta replicating plasmids can improve protein expression rate, due to higher copy number of the DNA material, and whether plasmid based expression is more susceptible to the effect of translesion DNA repair, compared to chromosomal DNA. Spores are formed during late stationary growth in nutrient deficient media. These exact conditions upregulate low-fidelity translesion polymerases and we hypothesized that eliminating these polymerases would improve expression homogeneity. Thus we additionally probe the effect of translesion polymerases by studying expression in two PolY, translesion polymerase, knockouts compared to wild-type *B. subtilis* (Fig 1B).

In this study we also designed and demonstrated engineered riboswitch regulation in *B. subtilis* using thermodynamic modeling to predict custom switch sequences for the control of red fluorescent protein expression^17^ (Fig 1C). This was done to create a protein expression sequence that would better report sequence changes due to a longer range of potential mutation sites (Fig 1D). This riboswitch, triggered by theophylline, also demonstrates the concept of an engineered, spore-based whole-cell biosensor and clearly illustrates the effect of a heterogeneous response and improvements caused by changes in plasmid type and cell line.

## Results

### Creating *B. subtilis* strains with riboswitch regulation of protein expression

Two of the most common methods for recombinant protein expression in *B. subtilis* rely on integrative or theta replicating vectors, with the majority of studies using the former due to the stated stability enhancement. In this study we compared a BioBrick 2.0 integrative plasmid, pBS1C, and a theta replicating pHT type plasmid^18^. A theophylline riboswitch-controlled RFP expression cassette was designed and cloned into both plasmids to develop an expression system that was especially sensitive to sequence fidelity (Fig. 1D); this is due to the increase in 5’ untranslated region (UTR) sequence length, giving a greater probability of point mutation occurrence. Along with the two types of expression vectors, three *B. subtilis* strains were studied: wild type strain (*B. subtilis* 168), a PolYA knockout of 168, and a PolYB knockout of 168^19,20^ (Fig. 1B). PolY knockouts were used in this study to evaluate the impact of error prone translesion DNA repair on expression homogeneity. Each of these cell lines were transformed with either theta replicating or integrating plasmids to yield a total of 6 expression strains for this study.

### Establishing stability of theta replicating, pHT, plasmids during vegetative growth and sporulation in *B. subtilis*

Protein expression from non-integrating plasmids is complicated by problems with structural stability^21^. Recently, a new type of theta replicating plasmids, pHT type, has been developed for *B. subtilis* to improve stability. pHT plasmids have been propagated in *E. coli* over 100 generations and demonstrated good retention with no recombination^16^. While sequence stability of a plasmid during propagation in *E. coli* is important, it does not guarantee stability during the vegetative growth or sporulation in *B. subtilis*. The stability of the expression template stored on a plasmid must be assessed, as time delayed protein expression relies on it being undamaged and usable upon germination. This has not been shown previously for theta replicating plasmids. Beyond that, plasmid retention during sporulation is highly specific to plasmid type and must be assessed empirically^22^. Stable chromosomal integration ensures retention of the genetic template throughout sporulation and thus is not evaluated here.

To evaluate the stability of a pHT plasmid during prolonged vegetative growth, cells were cultured for approximately 100 generations and harvested every 10 generations. Plasmid DNA was extracted from collected cell fractions and analyzed with agarose gel electrophoresis. Additionally, extracted plasmid material was used as a template in PCR reactions to amplify the protein expression region. We observed that the position of DNA bands of the plasmid and PCR product, 9116 and 1298 bases long respectively, is consistent in all lanes, and, therefore, preserved over all generations tested (Fig. 2A).

**Figure 2.**
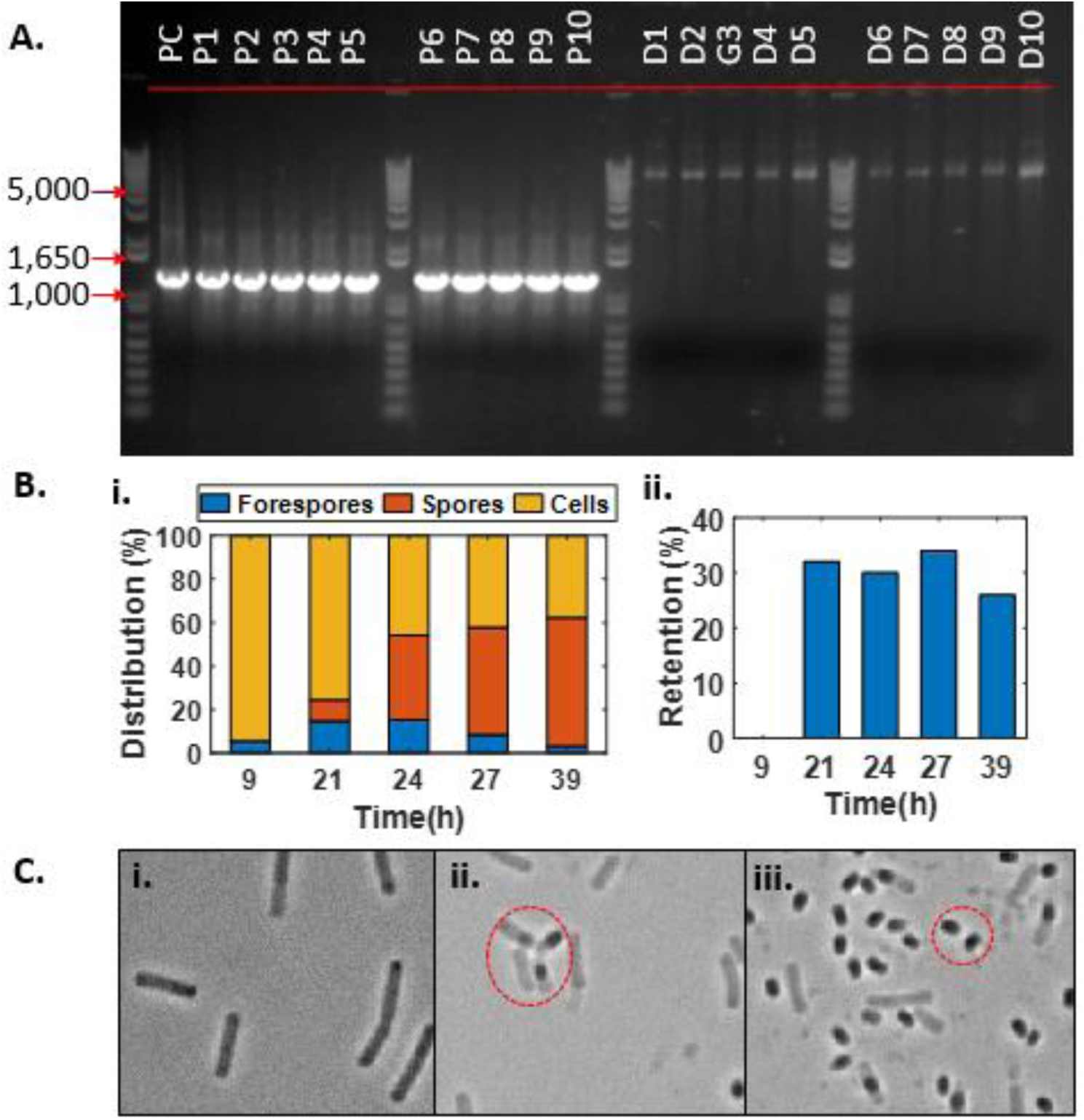
Plasmid stability in B. subtilis cells. (**A**) DNA gel electrophoresis results of Theta plasmid and PCR amplified expression region propagated in wild type *B. subtilis* cells. (**B**) Distributions of different *B. subtilis* cell phases during sporulation (i) and retention of plasmid (ii). (**C**) Examples of different cell types of B. subtilis: i. vegetative cells. ii. forespores. iii. spores.

To evaluate the stability of a pHT plasmid during sporulation, *B. subtilis* cells were grown in DIFCO sporulation media, samples were pulled at various growth phases – before the log growth phase, during early mid as well as late growth phases and in the stationary phase. Harvested fractions were assessed for plasmid retention via replica plating. The extent of sporulation was monitored using brightfield microscopy (Fig. 2C). Plasmid retention was measured when first spores were developed, 21 hours after the beginning of growth, and until the decrease in forespore concentration stabilized, at 39 hours. Plasmid retention was maintained throughout sporulation at 24-36% (Fig. 2B and 2C). These results indicate that about a quarter to a third of the spore population will express the target protein upon germination using the pHT plasmid, assuming that no subsequent horizontal gene transfer would take place that would further increase the population of cells expressing RFP.

### Quantifying heterogeneity of protein expression in *B. subtilis* mutants

The first step in determining the heterogeneity of protein expression was establishing a method to extract kinetic growth parameters of single variants. This was done by selecting colonies from each combination of cell, vector, and treatment condition (*e.g.* before sporulation, after sporulation) and growing them up individually on a plate reader. For each set of conditions, we had 30 replicates. Each growth was monitored for cell density (*via* absorption at 600 nm) and amount of protein present (*via* fluorescent signal). Both signals were converted to approximate, real units (cells/µl and µg/ml) using empirically derived transfer functions (Fig. 3A, Supplement 1). Moreover, these cell density and fluorescent data were fit with a 4 parameter logistics (4PL) and second order plus dead time (SOPDT) models to a good degree of statistical confidence (Supplement 1). These models allow us to reduce a large amount of growth and fluorescence data to analytical functions that can be evaluated to determine kinetic parameters as well as the lag time between cell growth and RFP expression.

**Figure 3.**
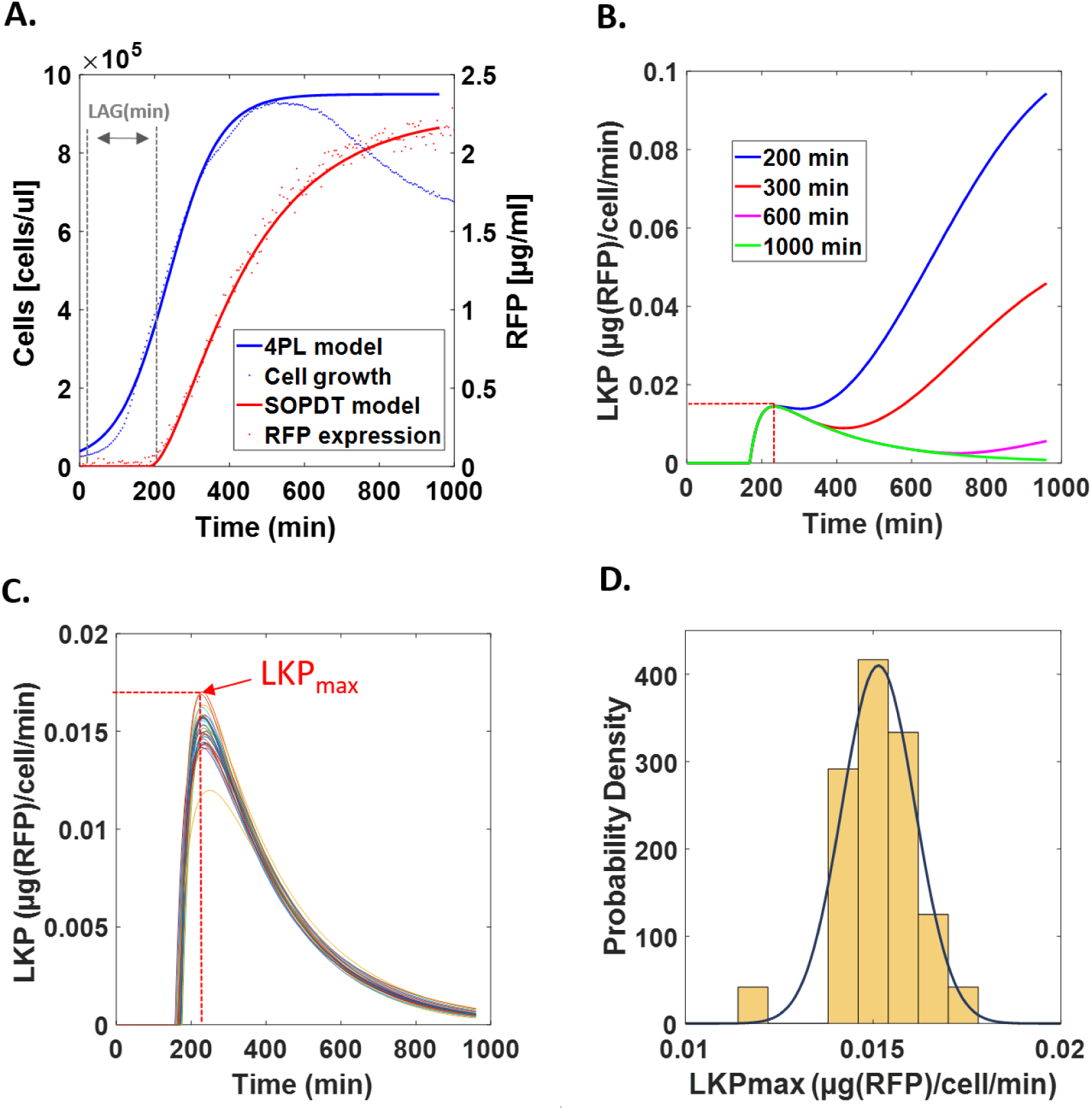
Raw empirical data and kinetic model fit. (**A**) Cell growth monitored via OD600 and fit with a four parameter logistics (4PL) model. RFP expression monitored via fluorescent signal and fit with a Second Order Plus Dead Time (SOPDT) model. Lag time between start of cell growth and start of protein expression is also noted on the raw data plots. (**B**) Lumped Kinetic Parameter (LKP) as a function of time evaluated from data in (**A**) using various viable cell durations. (**C**) LKP vs. time from colony replicates of same condition set used in (**A**) showing variations in maximum expression rates (LKP_MAX_). (**D**) Describing the variation observed in LKP_MAX_ from (**C**) using a normal probability density function.

A first principles, kinetic model was developed to extract kinetic parameters from the experimental data. We assume that a first order rate (k_1_) governs the transcription of DNA to RNA, and likewise RNA to folded protein (k_3_). Also we assume that RNA and protein also undergo degradation according to first order rates (k_2_ and k_4_ respectively). This reaction cascade can be mapped as displayed in Scheme 1.

> **Scheme 1** – Kinetic model

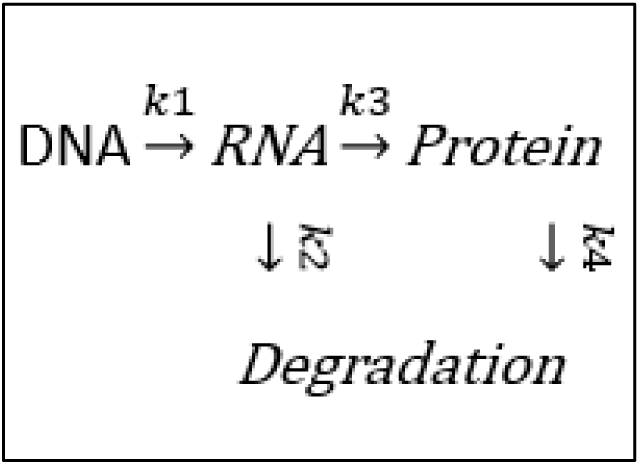

By assuming that 1) DNA concentration is proportional to the cell concentration (multiplied by a scalar c_1_), 2) RNA production is at steady state (no accumulation), and 3) the rate of protein degradation is negligible (k_4_ << k_2_) on the time scale of our experiments, we can express the rate of protein production with the following rate equation^23–25^:

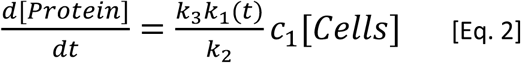

A few of these rate ‘constants’, especially the rate of mRNA transcription (k_1_) are known to be time variant^26,27^. The rate will change over the course of cell culture growth, depending on the promoter that is being used. A few careful studies have shown how each of these kinetic parameters can be isolated and empirically determined, however this is at the cost of low experimental throughput^26^. In this work, where we want to evaluate 30 independent clones for each cell-template-growth condition, such detailed analysis would be intractable. We are more interested in how the overall, or combined, kinetic rate changes amongst the different populations, thus for our work we report a lumped kinetic parameter (LKP) which again is time variant and has units of µg RFP produced per cell per minute (Eq. 3).

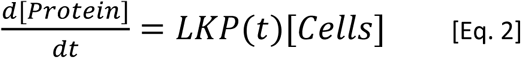

Using this equation, we can then solve for LKP as a function of time using the fit data models (Fig. 3B). This is done by determining the numerical derivative of the RFP expression curve at each time point and dividing by the cell concentration. One aspect not addressed by this model is the duration of viable cells. As written, it assumes that cells grow, contribute to the OD reading, and then produce protein for the entire experimental observation time. This is not the case; the cells will grow, produce protein for a time, and then die while still contributing to the OD signal. The solution of Eq.2 with the experimental data models can be adjusted to account for the cell viability time (Supplement 4), with various viability times modeled in Fig. 3B. For all cell viability times, the LKP peaks and then diverges. As the long term behavior of the models are more suspect anyway (note the drop in the experimental OD reading at long times, likely caused by biofilm growth and occlusion of the spectrophotometer light path), we focus our analysis solely on this peak LKP value (defined as LKP_max_). By pooling the clonal variants for a given condition set (cell type, vector type, and sporulation stage) we can see heterogeneity of the LKP_max_ value (Fig. 3C) which are described well by a normal distribution (Fig. 3D). We also determine the lag time between initiation of cell growth and expression of protein (displayed on Fig 3A) using the raw growth data and time delay model parameter from the SOPDT model fit to the protein expression data. These were pooled from different clonal variants and were also described well by a normal distribution.

### Characterizing expression heterogeneity for different plasmid and cell types

Using this method for extracting LKP_max_ values and lag times, we determined normal distributions of LKP_max_ and lag times for n=30 colonies for each cell and plasmid type (Fig. 4). We observe LKP_max_ values 12.58-fold higher for wild type *B. subtilis* cells that expressed RFP from the theta replicating plasmids compared to the chromosome integrated vectors (Fig. 4A). However, this seems to come at the expense of expression homogeneity as the variance of LKP_max_ values obtained for cells that express RFP from the theta replicating plasmid is 123 times that of cassettes integrated in the chromosome for the wild type strain (Fig. 4A and Supplement 1). The lag time distributions (Fig. 4B) show plasmid based expression in the wild type strain begins 60 minutes sooner than chromosomal based expression and has 15x less variance. It is important to note that these observations were not due to issues of lower signal to noise ratios leading to worse model fits and higher variance; the coefficient of variance for each experiment fit can be seen in Supplement 1.

**Figure 4.**
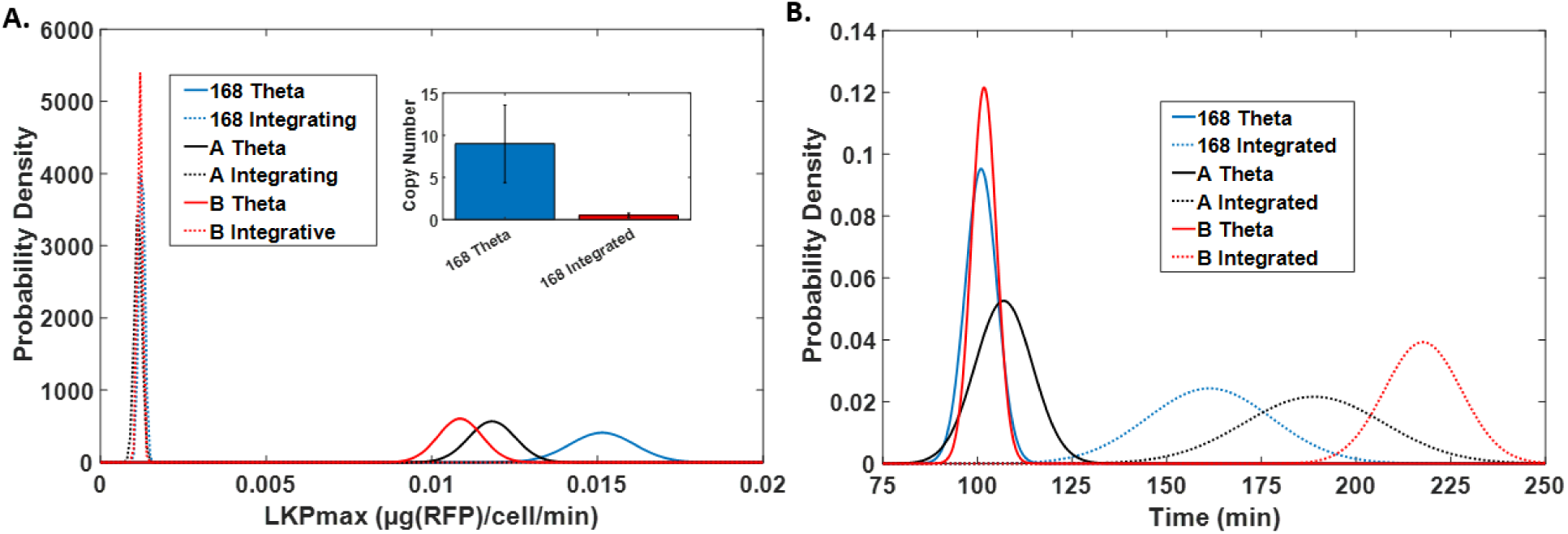
Comparing expression from a theta replicating plasmid and integrated DNA template. (**A**) Distributions of LKP_max_ values of *B. subtilis* strains 168, PolYA- (A), PolYB-with integrated and theta plasmid DNA. Insert shows relative copy number of expression fragments found in cells with either theta plasmid or integrated DNA. (**B**) Distributions of lag time values of *B. subtilis* strains 168, PolYA- (A), and PolYB-with either integrated or theta plasmid DNA. Each distribution is composed of n=30 colony samples.

The results acquired from this LKP analysis can also be visualized on a three dimensional plot to observe population variance and evaluate if the sample size would be sufficient to differentiate between cell and DNA types (Fig. 5). The majority of clustering can be seen along the LKP_max_ axis, which is also confirmed with principal component analysis showing 99.40% of variance is captured with a single principal component (Fig. 5 insert). Such clustering portends the ability to use machine learning to select clonal populations with desired expression rates, thus further improving the rate and homogeneity of the desired product.

**Figure 5.**
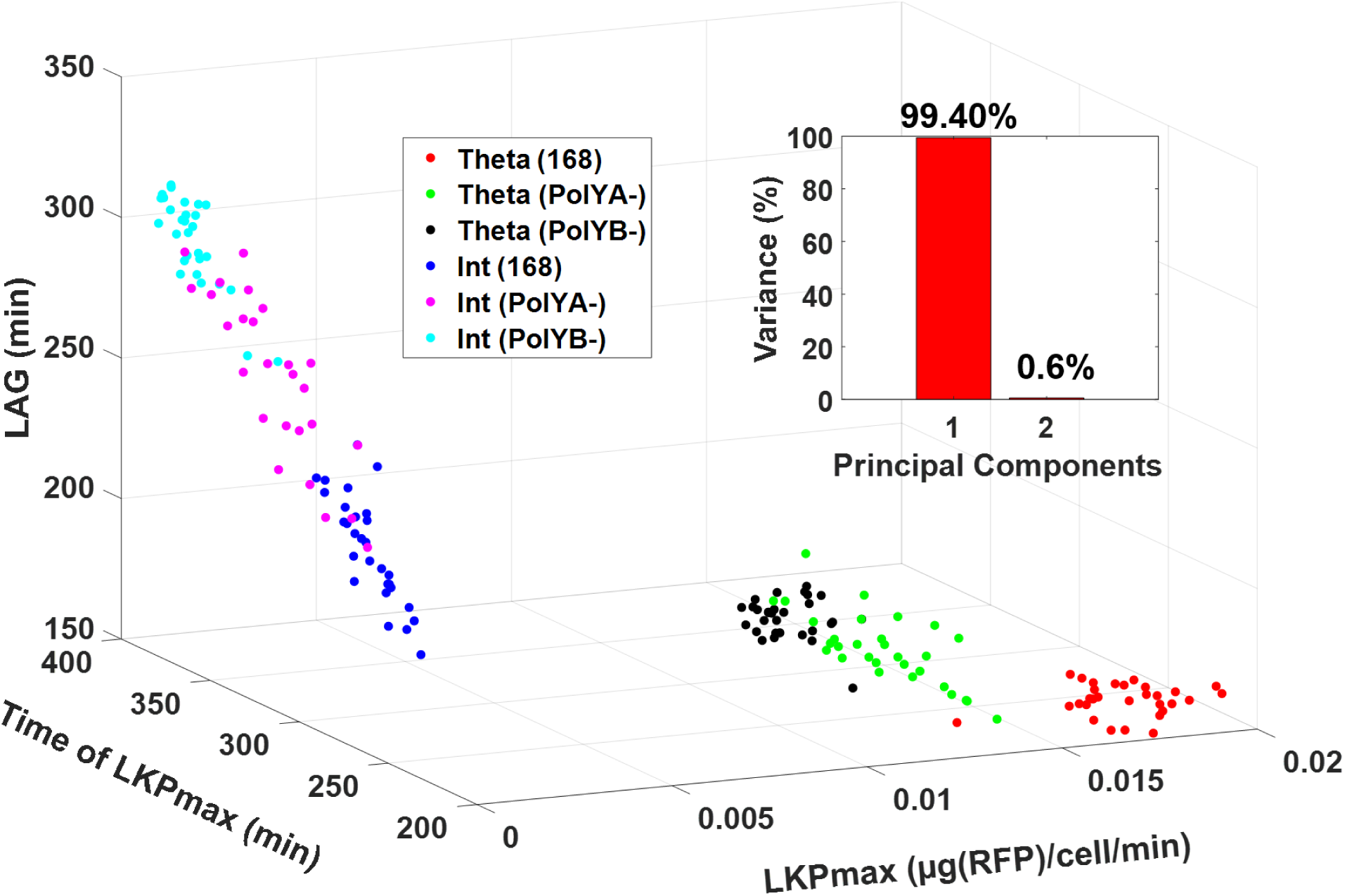
3D scatter plot representation of LKP model parameters from experiments used in Fig 4 (n = 30 for each cell and DNA combination). Insert shows principal component analysis of the LKP model parameters.

### Evaluating copy number and polymerase accessibility in plasmids and chromosomes

The heterogeneity observed may be due to differences in DNA copy number, polymerase accessibility for transcription, and segregational stability. Plasmids are often present as multiple copies inside of a bacterial cell and this may differ from the copy numbers observed from the integrating plasmids. The polymerase accessibility or level of transcription in a theta plasmid compared to a chromosome integrated template has not been well studied. To determine the role of plasmid copy number and transcription accessibility on these observations of heterogeneity, qPCR and RT-qPCR experiments were also performed to determine the concentrations of expression cassette DNA and mRNA, respectively.

*B. subtilis* cells were cultured in a microplate reader and fractions were harvested at beginning of growth, early, mid and late log, and stationary phases. Relative copy number of the expression cassette was compared to 16S rDNA (a normalizing standard) for each time point as previously described in the literature (supplement 5)^28^. Since the copy number was maintained at relatively constant levels throughout the growth phases, all time points were averaged (Fig. 4A insert, Supplement 5). The average relative copy number of the expression cassette on a theta plasmid was 9.4 relative copies and 0.6 relative copies for the integrated cassette, suggesting a plasmid copy number of approximately 15.

While differences in expression levels can be attributed to lower copy number of the template DNA in chromosomal expression, it is possible that the accessibility of template DNA also plays a role in affecting transcription levels. The concentration of expression cassette RNA was evaluated with RT-qPCR, and normalized by DNA copy number of the expression cassette (Fig 6A and 6B). The results show that DNA is more accessible (*i.e.* more RNA is produced per unit of DNA) when it is encoded in the chromosome, despite the fact that the promoter and 3’-UTR regions are identical. Therefore, the increased rate of protein production observed during expression from theta plasmid is caused primarily by DNA copy number.

**Figure 6.**
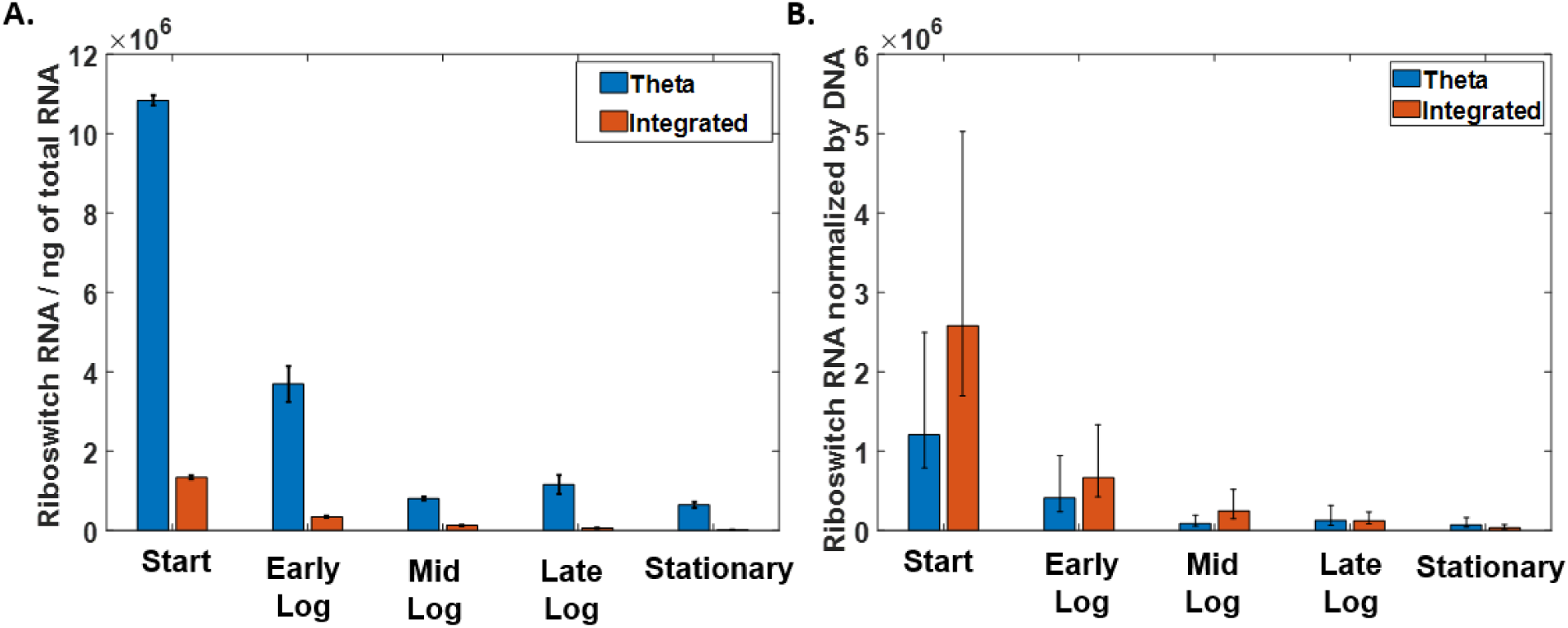
RT-qPCR of theophylline riboswitch RNA. **(A)** Copies of the riboswitch RNA normalized by total ng of RNA used in the RT-qPCR reaction. Showing one standard deviation from 2 replicates **(B)** Copies of the riboswitch RNA normalized by total ng of RNA and average copy number of copies of DNA in the cell. Showing propagated error (68% confidence interval) from RT-qPCR and qPCR experiments).

The distribution of lag parameters showed that noticeable expression of the target protein, RFP, begins sooner if the expression cassette is located on the plasmid rather than integrated in the chromosome. Revisiting that finding in the context of this qPCR and RT-qPCR data enables us to state that higher copy number of template DNA is the primary determinant of the yield and response time to the theophylline ligand. This is assuming that the riboswitch activation was not altered for either type of expression because the sequence was the same.

### Characterizing expression heterogeneity after sporulation using pHT plasmid

LKP_max_ and lag time distributions were then obtained from the wild type and mutant strains carrying a theta replicating plasmid before and after sporulation (Fig. 7). We observed that the variance of LKP_max_ values increased 2.07-fold after sporulation for the wild type strain. However, both PolY knockouts maintain tighter probability distributions compared to the wild type at the expense of slight (1.38-fold) reduction in protein expression rate. Nevertheless, this reduction is minimal since it is still 8.67 times higher than with DNA integrated in the chromosome.

**Figure 7.**
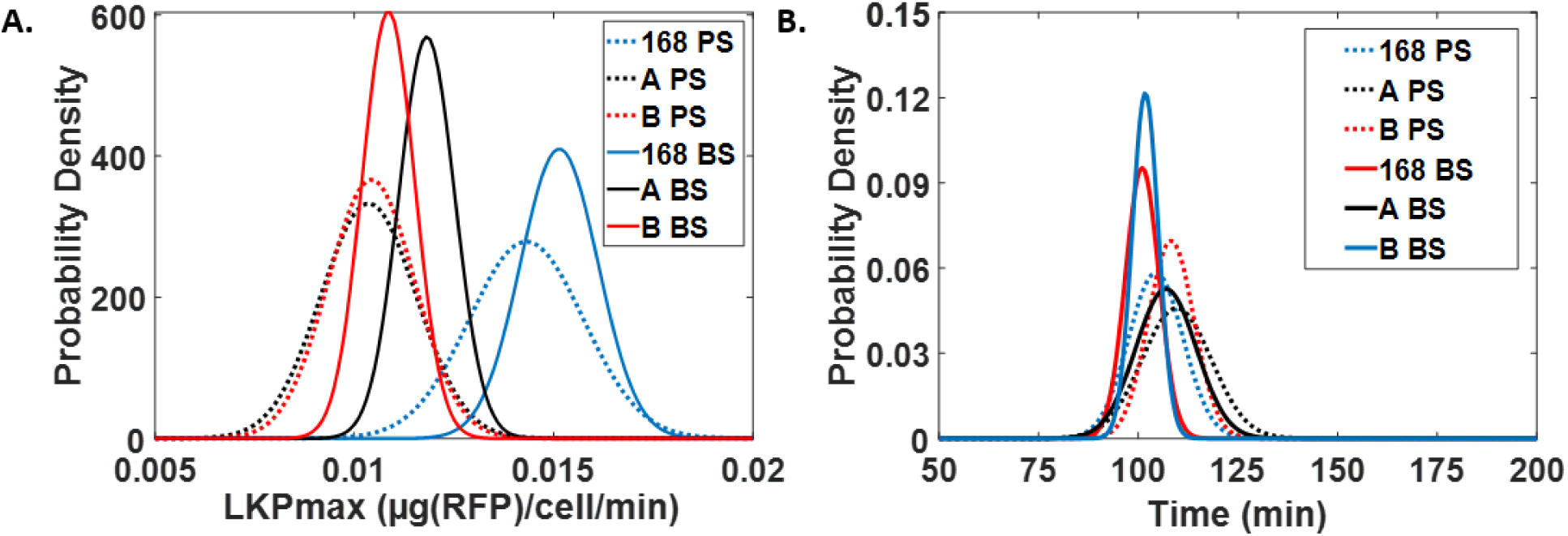
Distributions of LKP_max_ (**A**) and LAG times (**B**) before (BS) and post sporulation (PS) for strains 168, PolYB- (B), PolYA- (A) carrying a theta replicating plasmid.

In addition to studying the effect of sporulation on germinated cells in suspended cultures, we also performed surface, biofilm growth studies to observe the effect during adherent cell outgrowth. Images of bacterial biofilms formed before and after sporulation (n = 20 for each condition set) were examined using fluorescence microscopy (Fig. 8). Before sporulation the wild type cells produce more RFP compared to the PolyYB knockout. Both show dramatic reduction in expression after sporulation, however the PolyYB knockout exhibits higher levels of RFP post sporulation.

**Figure 8.**
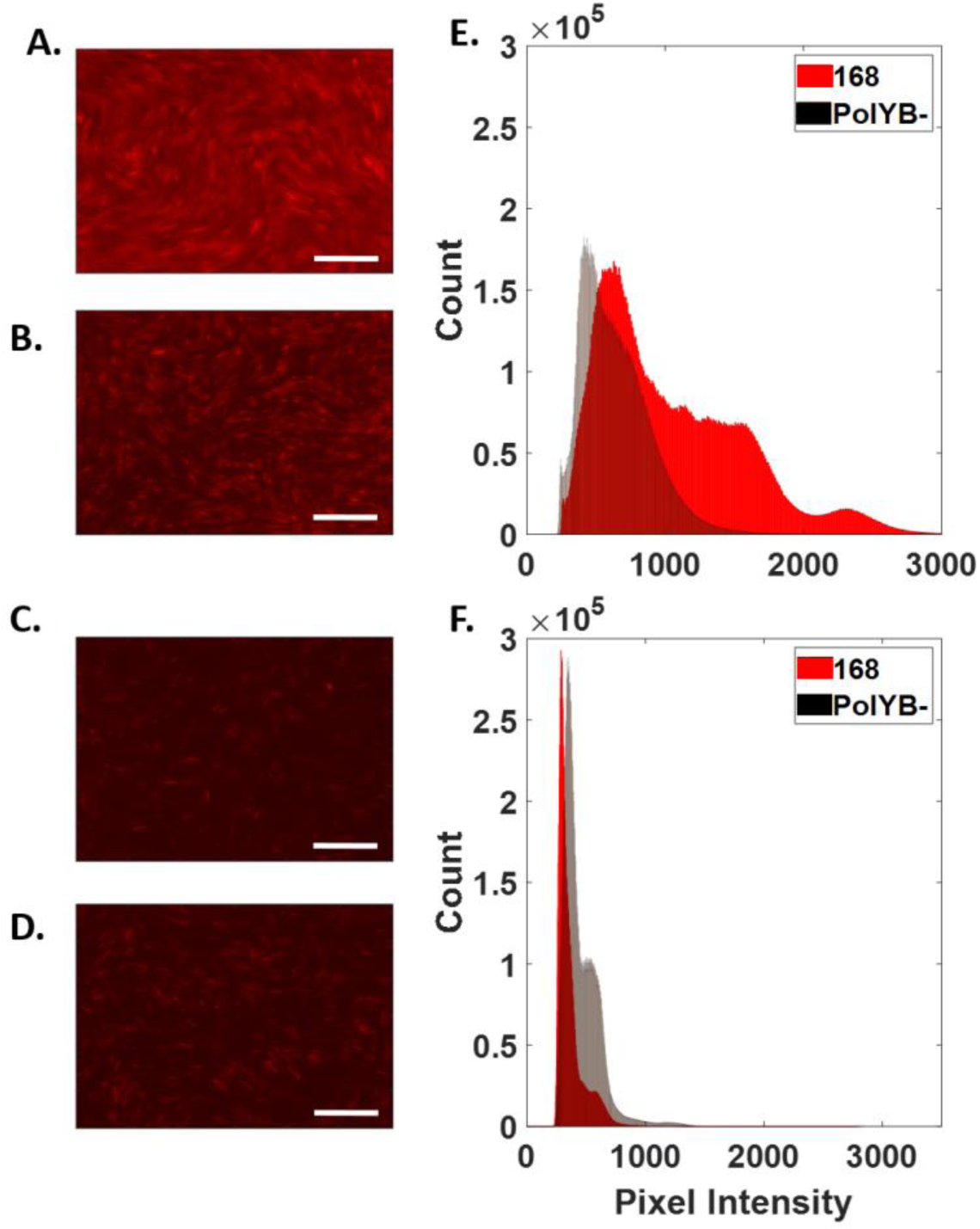
Microscopy image analysis of *B. subtilis* biofilms. All micrographs shown on same intensity scale (0 to 3500 RFU) to allow for comparison of intensity. Scale bar 20µm. (**A**) Biofilm of strain 168 before sporulation. (**B**) Biofilm of strain PolYB-before sporulation. (**C**) Biofilm of strain 168 after sporulation. (**D**) Biofilm of strain PolYB-after sporulation. Histograms of pixel intensities obtained from 20 images at each condition (A-D) showing general population behavior before (**E**) and after (**F**) sporulation.

### Comparing post-sporulation behavior of *B. subtilis* cells to UV damage behavior

During late stationary phase growth, bacteria exhibit low fidelity DNA replication due to translesion amplifications. A similar mechanism occurs during UV induced damage. In order to support our hypothesis that sporulation decreases DNA sequence fidelity, we also applied our LKP model to cells that were treated with varying UV exposures (7, 9, 11, 13, and 15 minutes). It is evident that longer UV treatment of cells generally increases variance of both the LKP_max_ and lag time distributions (Fig 9). It is also important to note that this effect on variance is much less pronounced in the PolYB knockout, as we already observed in the suspended and adherent cell tests. This data also supports the hypothesis that sporulation has a negative impact on DNA sequence fidelity because sporulated cells exhibit an increase in variance of LKP parameters similarly to UV treated cells.

**Figure 9.**
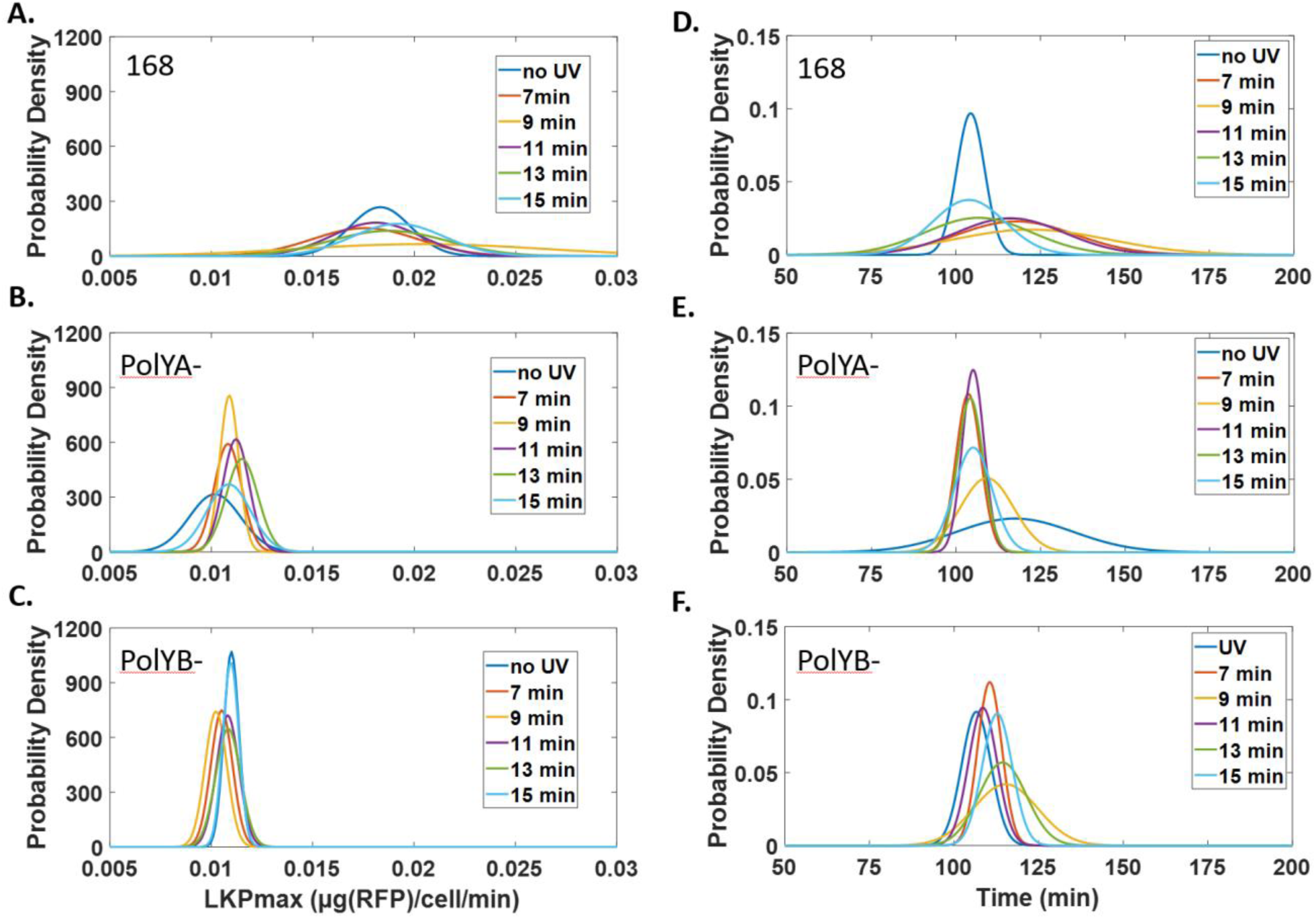
Evaluating effect of UV exposure on LKP_max_ and LAG time distributions for *B. subtilis* strains 168, PolYA- and PolYB- (n = 18 for each distribution). (**A**) LKP_max_ for strain 168. (**B**) LKP_max_ for strain PolYA- (**C**) LKP_max_ for strain PolYB-. (**D**) LAG time for strain 168. (**E**) LAG time for strain PolYA-. (**F**) LAG time for strain PolYB-.

## Discussion

With this work, we have shown that the backbone of the pHT-type theta replicating plasmid is resistant to homology driven recombination in *B. subtilis* during vegetative growth and sporulation. In addition, we have shown that this plasmid is present at higher copy numbers compared to DNA integrated into the genome (Fig. 6). However, since only about a quarter to a third of spores contain the plasmid upon germination (Fig. 2B), we hypothesize that these cells account for the increased copy number by carrying a plasmid that was concatemerized during transformation^29^. Given that homology driven recombination is possible in *B. subtilis*, it is unclear if the plasmid persists in a constant oligomeric state or whether it consistently changes the number of repeats present on the plasmid^30^. This might address higher error bars associated with Theta plasmid qPCR data (Fig. 4A insert). An alternative explanation might suggest localization of plasmids in *B. subtilis*, and stochastic release and incorporation into *B. subtilis* spores^31^.

Based on RT-qPCR data (Fig. 6B), we observe more RNA per strand of DNA for the integrated sequence *vs.* the theta-replicated construct. We attribute this to genomic DNA being more accessible to RNA polymerases in these cells. We therefore conclude that higher protein rates of expression observed in cells with theta replicating plasmids (Fig. 4A) are attributed primarily to the copy number of the expression region rather than the accessibility of DNA or rate of transcription.

We have demonstrated that spores of *B. subtilis* are good candidates for time delayed protein expression applications such as ‘on-demand’ biologics production, biosensors, or vaccine delivery applications. We bring attention to an issue that has been overlooked by other studies proposing the use of *B. subtilis* spores as an expression chassis: the fact that spores are generated in challenging media and during very late stages of growth, which is when low fidelity polymerases are most active, and therefore can introduce mutations to coding sequences of the encoded protein product. In this study we indirectly observe this effect through population changes in expression rates caused by point mutations to the riboswitch regulated production of a reporter protein. In follow up work, this effect can be confirmed through direct measure of protein and RNA sequences using mass spectrometry and next generation sequencing respectively. In the context of ‘on-demand’ therapies, such sequence heterogeneity would decrease efficacy and potentially increase immunogenicity. For whole cell biosensors these changes would affect the calibration curve, as varying lag times to expression and levels of expression would change this calibration. For vaccines, it is still unclear whether a lightly randomized distribution of produced antigen sequences would have a positive or negative effect on the quality of acquired immunity^32^. In each of these cases a decrease in expression variance is needed. This work has determined that the use of Y-family polymerase lacking mutants of *B. subtilis* are capable of decreasing the expression variance while having no effect on the bacteria’s ability to sporulate and whilst still maintaining a high level of expression using theta replicating plasmids.

This project emphasizes a tradeoff between expression rate and homogeneity from germinated *B. subtilis* spores. The faster rates with shorter lag times are the cells that have the largest variance. This observation enables judicious choice in selecting cell and template combinations based on intended applications. For example, faster expression would be desired for whole-cell biosensor applications but the variance would have to be accounted in the sensor model; for biopharmaceutical production, slower rates of expression may be tolerated thus ensuring a more homogenous product. In the context of oral vaccination, higher protein expression yields might be desired as they would minimize the number of doses required to generate effective immunity and the heterogeneity of the produced antigen may improve efficacy. In summary, time-delayed production of proteins from spores has great potential but care must be given in the choice of cell line and type of genetic template to match the desired production speed and tolerated variance.

## Methods

### LKP kinetic model of protein expression

The kinetic model and general approach is summarized in the results section. Cell density was measured using absorbance at 600 nm in BioTek Synergy Neo 2 plate reader, and fit with a 4 parameter logistics model, Eq.3. In this equation, t is time, d and a are maximum and minimum OD600 value reached by the cells respectively. Variable c is the point of inflection, while b, is the Hill’s slope of the curve at point c. Protein expression was measured as a function of fluorescence, and found to fit well to a Second Order Plus Dead Time (SOPDT) model, Eq. 4. In this equation, t is time, K is the gain, theta is the delay between data collection began and first detectable increase in RFU. The variables Tau_1_ and Tau_2_ are time constants that account for the curvature of the response. Protein and cell concentrations was measured directly using the plate reader.

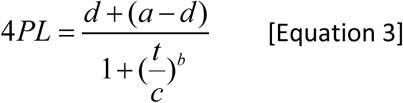

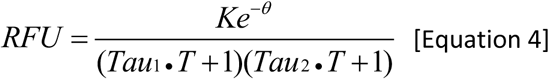

The lag time was calculated as the difference between the onset of cell growth and onset of protein expression. The onset of cell growth was evaluated as OD600 absorbance increases by 2 standard deviation values above the start baseline. The onset of protein expression was found by the theta parameter in the SOPDT model fit to the protein fluorescence data.

### DNA sequences and mutants

A computational model of riboswitch activation, the Riboswitch Calculator, was previously used to design theophylline-binding, translation-regulating riboswitches controlling the expression of mRFP1, or an mRFP1 fusion protein, in *E. coli*^33^ (Supplement 7). Given the sequence, secondary structure, and ligand-binding free energy of a theophylline RNA aptamer, the model designed pre- and post-aptamer sequences to minimize the mRNA’s translation rate in the OFF state (without ligand) and maximize the mRNA’s translation rate in the ON state (with ligand).

The theophylline riboswitch sequence was cloned into the pHT43 vector between restriction sites KpnI and AatII, and into pBS1C vector between cloning sites EcoRI and SpeI. Cloned template was propagated in Top10 *E. coli* cells. Cloning was confirmed with Sanger sequencing.

Each plasmid was transformed into either the wild type strain of *B. subtilis*, 168, or one of the mutants, which were the wild type strain with knocked out PolyA or PolYB genes^19,20^.

### Cell growth and protein expression

*B. subtilis* cells were transformed with either pHT or pBS1C plasmid containing the theophylline riboswitch using conventional methods ^34^. Positive transformants were selected on chloramphenicol (5 µg/ml) plates. Transformants were verified with Sanger sequencing, and stored as a glycerol stock with 20% glycerol concentration. If cells were used in the sporulation experiment, prior to plating, they were cultured for 5 days in DIFCO sporulation media. Then, they were plated on Chloramphenicol containing agarose plates.

Prior to every growth experiment, glycerol stocks were streaked on agarose plates to acquire individual colonies. Agarose plates were subjected to UV treatment for various amounts of time by incubation on the DNA transilluminator for the UV treatment experiment. Individual colonies were picked and grown overnight in a 5ml LB culture with chloramphenicol. Overnight cultures were diluted 1:10 into a 96-well microtiter plate. Plates were cultured for 24 hours at 37C, with maximum agitation in the BioTek Synergy neo 2 plate reader. Absorbance at 600nm was recorded as a measure of cellular density, and fluorescence with excitation and emission at 584nm and 607nm respectively as a measure of RFP concentration.

### RFP standard curve

In order to correlate RFU signal obtained from the plate reader with approximate protein concentration, protein standards were acquired by cloning the RFP sequence used in all of the experiments in a pMal vector and expressed in BL21 *E. coli* cells. The protein was purified in two steps using amylose and Ni-NTA affinity chromatography close to homogeneity. Protein concentration was assayed using Pierce 660 assay (Sigma), with BSA for standard. After determining the concentration of RFP in the solution, a serial dilution with fresh LB media was performed and protein concentration was correlated with fluorescence (Supplement 2).

### Stability of Theta replicating plasmids

To test the stability of the theta replicating plasmid, pHT43, *B. subtilis* cells were grown to saturation and diluted one thousand times. The procedure was repeated 10 times. The number of generation that existed during one growth period was estimated to be about 100. The entire experiment evaluated the stability of DNA over about 1,000 generations. 8 ml of cells were harvested, for plasmid extraction. The extract was used to amplify the gene of interest. Plasmid extract and the amplified region were stored at −20°C until gel electrophoresis. Both, the plasmid extract and the amplified gene were used for DNA gel electrophoresis (figure 2A).

### Plasmid retention

*B. subtilis* 168 cells carrying a theta plasmid were grown overnight in DIFCO sporulation media. The overnight was used to infect 250 ml of the same DIFCO sporulation media and grown for 3 days. Cells fractions were harvested at various time points throughout the growth. These fractions were imaged using brightfield microscopy to verify the distribution of vegetative cells to forespores to spores.

Harvested fractions were washed in double deionized water twice. Lysozyme was added to washed cell mixture and incubated at room temperature for 30 min. Then, bacterial protease was added to the mixture and the sample was incubated at room temperature for 30 min. This treatment destroyed all vegetative cells and forespores. After that, a serial dilution was performed to identify an optimal concentration of cells, and plated on LB agarose plates that did not contain antibiotics. 100 Individual colonies were picked and streaked on agarose plates containing chloramphenicol. The number of colonies that grew on chloramphenicol plates was divided by 100. The result represented the fraction of cells that retained the plasmid.

### qPCR and RT-qPCR

*B. subtilis* cells strains 168 with either a pHT or pBS1C plasmids were grown in the microplate reader in a 6 well plate. The first 5 wells were harvested about every 2 hours from the beginning of growth. Time points 1 through 5 represent stages of growth that were spaced out every 2 hours starting from infection. Cells were then diluted until the final absorbance at 600 nm was 10 or below depending on the total cell mass that was available. Cells were lysed by 15 min pretreatment with lysozyme followed by bead beating. Samples were stored in −80°C freezer until they were used for qPCR. No DNA purification was performed for qPCR reactions, instead unpurified cell lysate was used. RNA was extracted with Qiagen RNeasy kit. The reactions were setup with NEB’s Luna Probe qPCR or RT-qPCR master mix with 0.5 µl of template in 20 µl reactions. Reactions were performed in the ABI StepOne plus thermocycler.

Data was processed according to MIQE guidelines and fit with a 5 parameter sigmoidal curve using custom scripts written in Matlab ^35^ (Supplement 5).

Lysed cell fractions were serially diluted 10 fold five times and 0.5 µl of template was used in 20 µl reactions in duplicates. Same cell lysate fractions were used for both the reference and riboswitch probes. Three of those dilutions were used with a TaqMan probe designed for the theophylline riboswitch sequence. Relative difference in concentrations between the copy number of the theophylline riboswitch and the reference sequence were measured using the *2*^-Δ*Ct*^ formula and efficiency was calculated using the 10^(−1/*Slope*)^ −1 equation as previously described^35^.

Concentration of RNA that coded for the theophylline riboswitch was calculated using the absolute standard curve method. Standard was purified theta replicating plasmid that was propagated in *E. coli*. C_t_ values of the RNA were converted to copy number of theta plasmid. That value was then normalized by the total amount of RNA in the template used in the RT-qPCR reaction. As a result, the output of the experiment was relative RNA copy number per ng of extracted RNA. In order to estimate accessibility of RNA, the relative concentration of RNA per ng of extracted RNA was normalized by the average copy number of the plasmid from the qPCR experiment.

### Flow Cytometry

In order to convert OD600 nm values acquired via the plate reader to individual cell counts, a standard curve was created. First, a serial dilution of cells grown in the plate reader was generated. These cell concentrations were measured using the Spec20 spectrophotometer after diluting cell samples to fit in the OD600 nm 0.1-0.6 range. This generated a standard curve of OD600 nm acquired on the plate reader and real OD600 nm values. Then, cell fractions of OD600 values 0.4, 0.6, 1 and 1.4 were selected for flow cytometry experiments. Cells were diluted to the appropriate density using LB with 5% sodium azide solution. Then, cell samples were used with the BacLight bacterial viability and Counting Kit (Thermo Fisher). All flow cytometry experiments were performed on the factory-direct unmodified BD FACSCanto flow cytometer (San Jose, CA). We were using 488 nm laser for extinction, and detected fluorescence with a 525/550 nm as well as 610/620 nm bandpass filters.

## Acknowledgments

We thank Shawn Rigby, manager of the flow cytometry facility at Iowa State, for assisting with conducting cell counting experiments. We thank Alexandra Scupham, USDA scientist, for consulting on qPCR theory and helping with data processing. We thank Daniel Zeigler, manager of BGSC, for helpful discussions and supplying all necessary Bacillus Subtilis strains and plasmids. Funding was provided by a Roy J. Carver Charitable Trust Seed Grant and ISU internal funds.

