## Supplement for "Controlling Heterogeneity and Increasing Titer from Riboswitch-Regulated *Bacillus subtilis* Spores for Time-Delayed Protein Expression Applications"

### 1. SOPDT and 4PL fits.

Cellular growth as a measure of OD at 600 nm was recorded and fitted with a 4 parameter logistics (4PL) curve. Expression of RFP was measured as a function of fluorescence of RFP 584/607 nm and fitted with a second order plus dead time model (SOPDT). Fit parameters were recorded for (1) theta vs integrating vector (2) UV treatment and (3) sporulation experiments (attached <modelParam.xlsx>).

### 2. Protein concentration standard curve:

In order to convert RFU data to meaningful units, we created a standard curve using Pierce660 assay. In this assay, a BSA standard protein solution (sigma) was diluted according to Supplement Table 1. The protein sample was mixed with the Pierce660 assay master mix according to the manufacturer’s protocol. Absorbance values at 660nm were measured and correlated with protein concentration (Supplement Figure 1). In order to correlate protein concentration with fluorescence, RFP was purified as described in the methods section, its fluorescence as well as absorbance at 660 nm were measured (<modelParam.xlsx> and Supplement Figure 2, Supplement Table 2).

Supplement Table 1 Dilution Table of BSA standard used in Pierce660 assay (supplement figure 1)

| **Sample** | **Concentration (mg/ml)** | **AVG A660nm** | **A660nm** | | |
| --- | --- | --- | --- | --- | --- |
| 1 | 1 | 0.456333333 | 0.445 | 0.462 | 0.462 |
| 2 | 0.5 | 0.336666667 | 0.34 | 0.331 | 0.339 |
| 3 | 0.25 | 0.272 | 0.269 | 0.274 | 0.273 |
| 4 | 0.125 | 0.237333333 | 0.232 | 0.238 | 0.242 |
| 5 | 0.0625 | 0.217 | 0.222 | 0.213 | 0.216 |
| 6 | 0.03125 | 0.203 | 0.204 | 0.197 | 0.208 |
| 7 | 0.015625 | 0.197 | 0.201 | 0.197 | 0.193 |
| 8 | 0.0078125 | 0.195 | 0.198 | 0.194 | 0.193 |
| 9 | 0.00390625 | 0.191333333 | 0.197 | 0.189 | 0.188 |
| 10 | 0.001953125 | 0.193333333 | 0.197 | 0.194 | 0.189 |
| unknown | | 0.256 | 0.257 | 0.256 | 0.255 |

Supplement Table 2 Fluorescence and corresponding protein concentration values of RFP used in Pierce660 assay (Supplement figure 2).

| **Concentration (mg/ml)** | **AVG RFU** | **RFU** | |
| --- | --- | --- | --- |
| 0.051881303 | 12041.5 | 12481 | 11602 |
| 0.025940652 | 5648.5 | 5744 | 5553 |
| 0.012970326 | 2684 | 2750 | 2618 |
| 0.006485163 | 1327 | 1331 | 1323 |
| 0.003242581 | 593.5 | 615 | 572 |
| 0.001621291 | 279.5 | 279 | 280 |
| 0.001621291 | 269.5 | 273 | 266 |
| 0 | 28 | 30 | 26 |

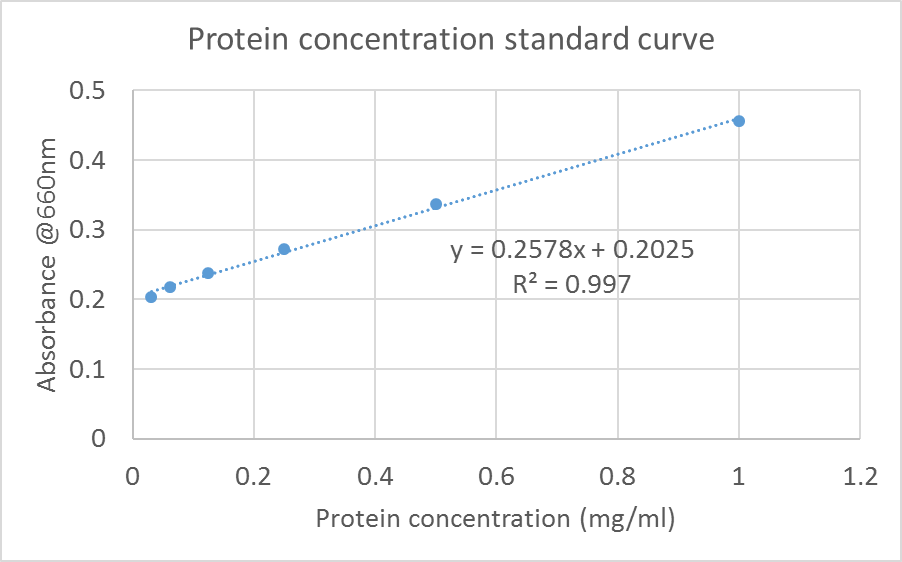

Supplement figure 1 Protein concentration and absorbance at 660nm standard curve.

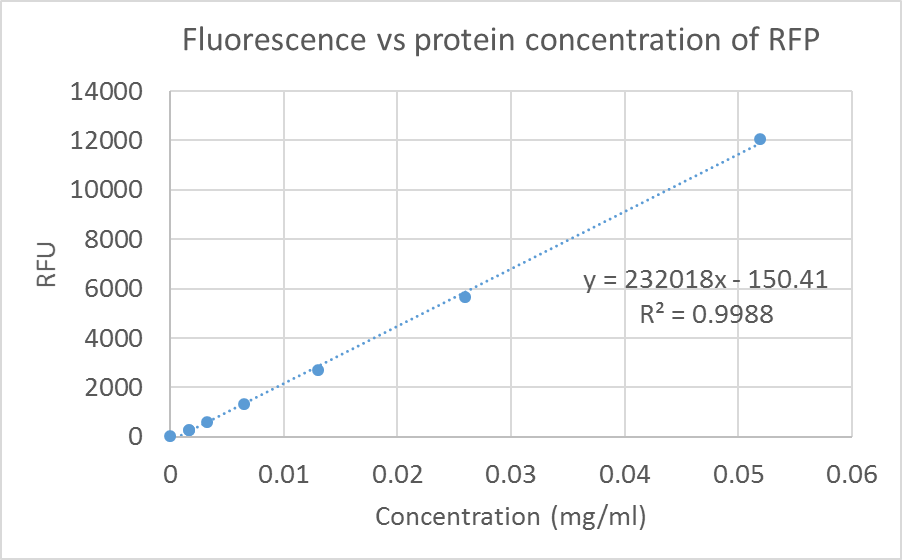

Supplement figure 2 Fluorescence and protein concentration of RFP standard curve.

### 3. Cell concentration standard curve.

B. subtilis cells were cultured in the plate reader, and cellular densities were approximated using absorbance at 600nm. In order to convert the absorbance values to meaningful concentration— cell count, flow cytometry was used. Cell samples were prepared as described in the methods section. First, absorbance at 600nm in the plate reader was converted to absorbance values with a constant 1cm path length (Supplement figure 3). Then, B. subtilis cell culture was diluted to concentrations that corresponded with OD600 values of 0.004 through 0.014 to be used in a flow cytometry experiment. A standard curve of cell count to optical density at 600 nm with path length of 1cm was created (supplement figure 4).

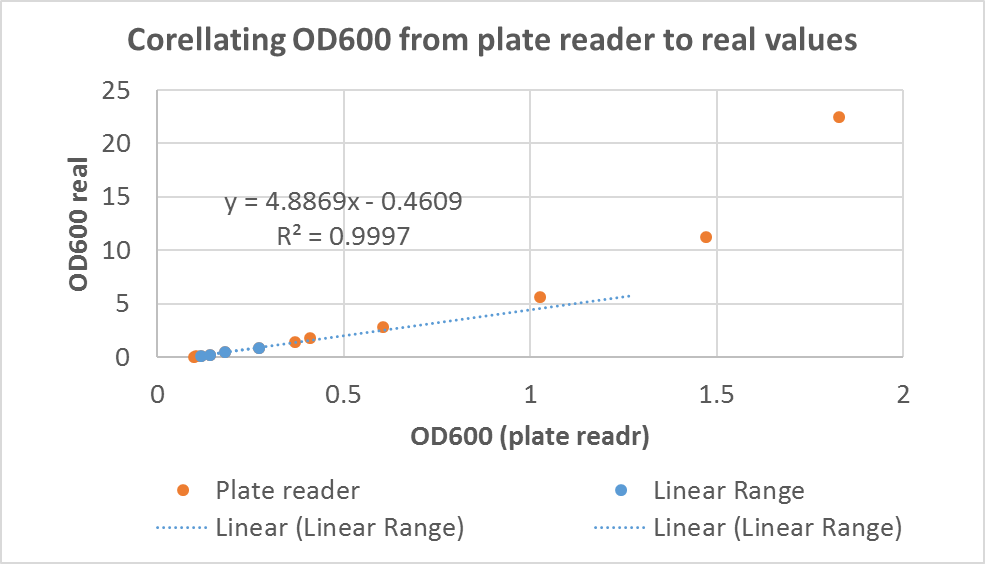

Supplement figure 3 Standard curve used to convert OD600 values acquired in the plate reader to 1 cm path length.

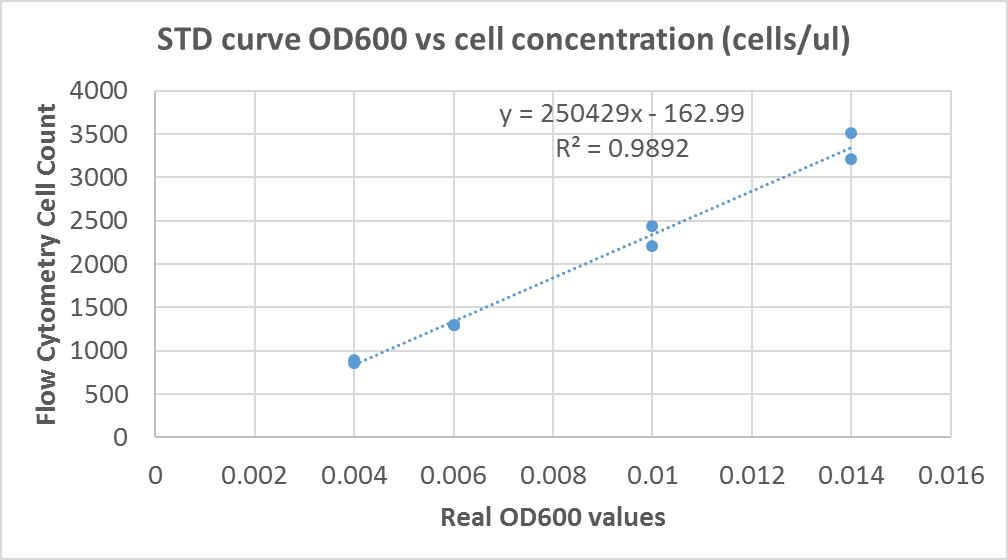

Supplement figure 4 Standard curved used to correlate cell counts and OD600 values

### 4. Matlab code for fitting 4PL and SOPDT models to raw data:

Below is a sample code that was used to reduce absorbance and fluorescence data to 4PL and SOPDT models, as well as extract LKPmax and LAG model parameters. This code was also used to generate normal distribution plots from LKPmax and LAG.

```
%Coded by DDT on 03/01/2019
%This is a sample of a code that was used to import raw OD600 and RFU data
%and fit it with 4PL and SOPDT respectively

function fitRawData
clc
clear
close all

%imports 3 types of data: time, OD600, F;
OD=xlsread('fileName.xlsx');
F=xlsread('fileName.xlsx');
time=xlsread('fileName.xlsx');
time=time*24*60; %convert to min

%Imports the row that contains names of wells
[~,posOD]=xlsread('fileName.xlsx');
[~,posF]=xlsread('fileName.xlsx');

%%
%Convert OD and RFU data to mg/ml and cells/ul units.
%used standard curves

ODreal=(4.8869.*OD)-0.4609; %convert to ODreal
cellCount=(250429.*ODreal)-162.99; %convert to Cell Counts

RFU2MET=((4E-6).*F)+0.0008; %mg/ml
RFU2MET=RFU2MET.*1000; %ug/ml

%%
%sort imported data
%The array below dictates the order by which to sort samples.
T={
    'B2' 'B3'    'B4'    'B5'    'B6' 'B7' 'B8' 'B9' 'B10' 'BTab' 'C2'   'C3'    'C4'    'C5'    'C6' 'C7' 'C8' 'C9' 'C10' 'C11'   'D2'  'D3'    'D4'    'D5'    'D6' 'D7' 'D8' 'D9' 'D10' 'D11'  'E2'   'E3'    'E4'    'E5'    'E6' 'E7' 'E8' 'E9' 'E10' 'E11'    'F2' 'F3'    'F4'    'F5'    'F6' 'F7' 'F8' 'F9' 'F10' 'F11'    'G2' 'G3'    'G4'    'G5'    'G6' 'G7' 'G8' 'G9' 'G10' 'G11';
};

all_OD=zeros(length(time),length(posOD));
all_F=zeros(length(time),length(posF));

%sorts OD data by column
for i=1:length(T)

   if find(strcmp(posOD,T(i))) ~=0
       id=find(strcmp(posOD,T(i)));
       all_OD(:,i)=cellCount(:,id);                                %%%
   end
end

%sorts F data by column
for i=1:length(T)

   if find(strcmp(posF,T(i))) ~=0
       id=find(strcmp(posF,T(i)));
   all_F(:,i)=RFU2MET(:,id);
   end
end

OD_model=[];
F_model=[];
modelParam = zeros(length(posOD),21); %21 parameters will be saved to the excell file


for i=1:1 %just run once because you are trying to get an example graph


    %id max OD600 values
    [~,max_OD]=max(all_OD(:,i));
    X=time(1:max_OD+20);         %increased to max_OD+20 to get real approximation of max OD in the fit
    Y=all_OD(1:max_OD+20,i);

    %Fit OD600 data
    % Fit type is a 4 parameter logistics curve, where a is min, d is max, c= time of inflection, b = slope at inflection
    try
    FT = fittype('d+(a-d)/(1+(x/c)^b)');
    [p,gof] = fit(X,Y,FT,'StartPoint',[3.046e+04 3.377 233.8 9.771e+05]);
    catch
        p.a = 0; p.b = 0; p.c = 0; p.d = 0; gof.rsquare = 0;
    end

    %regenerate the graph to see quality of fitting
    OD_model(:,i)= p.d+(p.a-p.d)./(1+(time./p.c).^p.b);

    %save model parameters
    modelParam(i,1:4)=[p.a p.b p.c p.d];
    modelParam(i,5)=gof.rsquare;


            %Fit F data
            %Fit type is SOPDT
            Y = all_F(:,i);
            X = time;
            try
            FT = fittype('K*(1-(tau1*exp(-(x-theta)/tau1)-tau2*exp(-(x-theta)/tau2))/(tau1-tau2))*heaviside(x-theta)+Rbak');
            [p5,gof5] = fit(X,Y,FT,'StartPoint',[500 70 160 114 222]);
            catch
                 p5.K = 0; p5.tau1 = 0; p5.tau2 = 0; p5.theta = 0; p5.Rbak = 0; gof5.rsquare = 0;
            end

            %regenerate the flurescence data to eval quality of the fit
            F_model(:,i,2) = p5.K*(1-(p5.tau1.*exp(-(time-p5.theta)./p5.tau1)-p5.tau2.*exp(-(time-p5.theta)./p5.tau2))./(p5.tau1-p5.tau2)).*heaviside(time-p5.theta)+p5.Rbak;

            %save fit parameters
            modelParam(i,11:15) = [p5.K p5.tau1 p5.tau2 p5.theta p5.Rbak];
            modelParam(i,16) = gof5.rsquare;


% What if we assume they can produce for a certain time period, and
% then they become dormant (that is, still showing up in the OD count,
% but not contributing to new RFP production)

t_persist = 1000; %minutes
tstep = tvec(3)-tvec(2); % time between steps in recorded data
nstep = round(t_persist/tstep);
ProductiveCells = movsum(CellNew,[nstep-1,0]); % This sums all of the 'NEW' cells in the time window specified for cell production
LKP_pc = Prate./ProductiveCells;

%calculate LKPmax
LKPmax{i,ii}=max(LKP_pc);

%now generate normal distributions
mu1=table2array(LKPmax(:,1));
mu1=mu1(mu1>0);
sig1=std(mu1);
mu1=mean(mu1);

f1 = exp(-(y-mu1).^2./(2*sig1^2))./(sig1*sqrt(2*pi)); %normal distr

mu1=table2array(LAG(:,1));
mu1=mu1(mu1>0);
sig1=std(mu1);
mu1=mean(mu1);

f1 = exp(-(y-mu1).^2./(2*sig1^2))./(sig1*sqrt(2*pi)); %normal distr

%Plot Figures

end
end
```

### 5. qPCR and RT-qPCR data processing.

All qPCR and RT-qPCR data was acquired on ABI StepOne Plus thermocycler. Both probes were cycled simultaneously with a 60 second initial denaturation at 95C, and then 40 cycles of 15 second denaturation at 95C followed by 30 second extension step at 60C. RT-qPCR samples were subjected to an additional reverse transcription step at 55C for 10 minutes.

Fluorescence data that was collected by the thermocycler was exported into an excel file. That data was then sorted using custom matlab code (qPCR_data_analyzer.m). Organized data was then fit with a 5 parameter logistics curve (qPCR2mat_V2.m). Raw data was normalized by subtracting fluorescence detected at the first step from fluorescence at each step of amplification. Normalized data was then converted to the log scale. Threshold values were then set manually. Linear standard curves were fitted through the Ct vs DNA or cell concentration data, in order to obtain the efficiency values (plotqPCR.m). When the efficiency values were confirmed to be in the same range, Ct values of the reference and unknowns were used to calculate relative differences in concentration using 2^-ΔC^_T_ method (Supplement figure 5).

This series of 3 programs was used to process qPCR data. In order to acquire samples for qPCR, cells, B. subtilis strain 168 with either a theta replicating or integrating plasmid were grown in a 6 well plate and harvested every 2 hours according to the methods section. Experiments were performed in duplicates, and 2 trials of qPCR was done on B. subtilis 168 with a theta plasmid and 2 trials on cells with the integrating plasmid (Supplement figures 6 - 9).

No growth phase dependent trends were observed, and, therefore, copy number measurements at all time points were average (Supplement figures 10 and 11). Since the reference, 16s rDNA, is present in B. subtilis cells in 10 copies, the 1:1 ratio of the theta plasmid was converted to 10 copies of the expression cassette. Conversely, since about a 1:10 ratio of reference 16s rDNA to integrated DNA was observed, the copy number of expression cassette integration was estimated to be equal to 1.

Lysed cell samples that were used in qPCR were stored in -80C and then thawed for RNA purification that was used the RT-qPCR experiment. RNA was extracted and purified according to the protocol described in the methods section. RNA concentrations were estimated using a ThermoFisher nanoDrop (Supplement Tables 3 - 4). RNA copy number was approximated using the absolute quantification method (Supplement figure 12). Theta plasmid with a theophylline riboswitch was propagated overnight in Top10 E. coli cells, and then miniprepped. An 8 point standard curve was prepared. The copy number of theophylline RNA was normalized by the total concentration of RNA in the RT-qPCR reaction (Supplement Table 5, figure 5A). Finally, specific copy number of RNA in the total pool of RNA was normalized by the copy number of DNA in order to compare the accessibility of the DNA template type (Supplement Table 6, figure 5B).

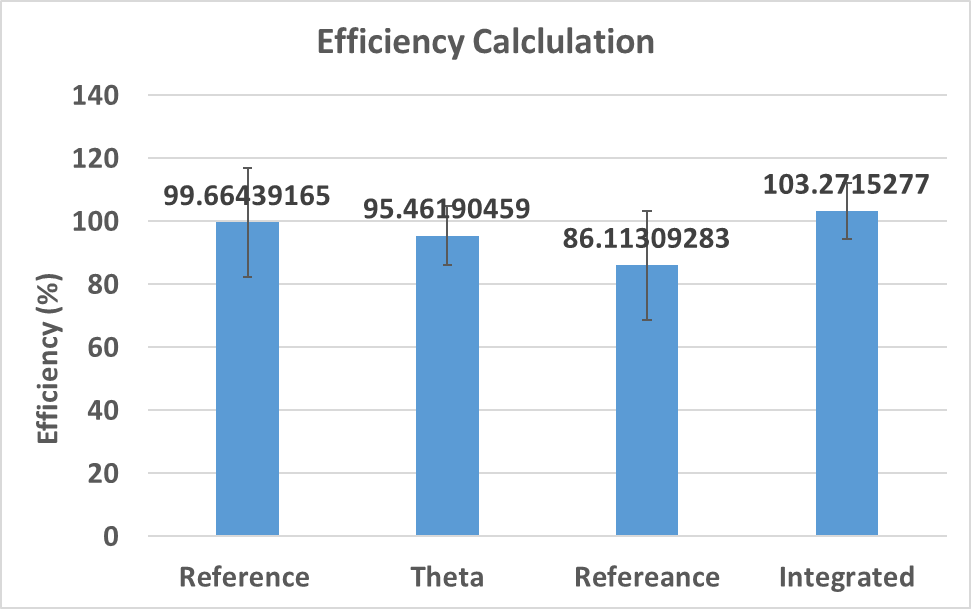

Supplement Figure 5 qPCR amplification efficiency.

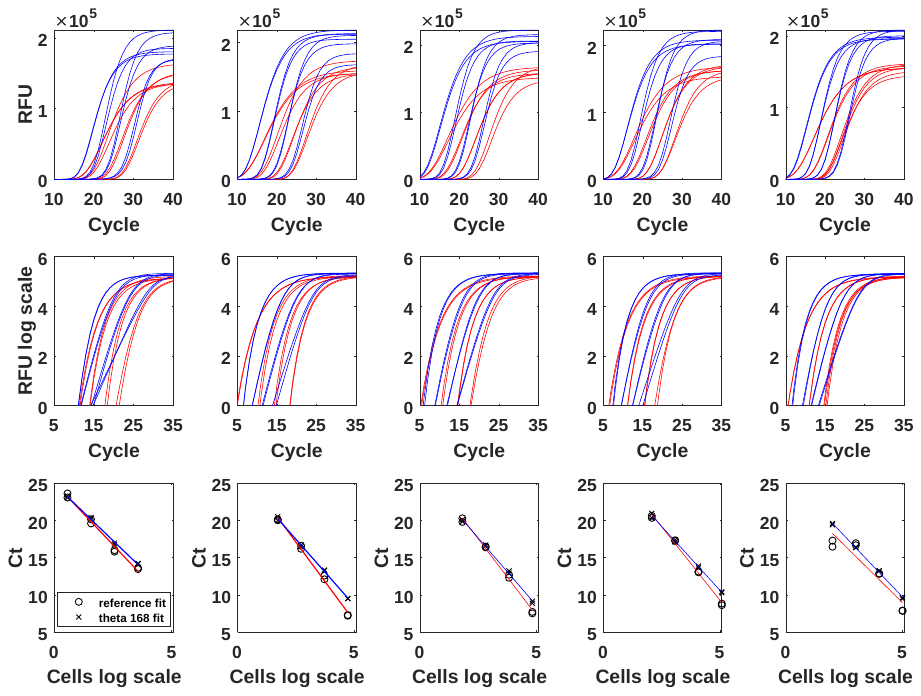

Supplement figure 6 qPCR data from B. subtilis cells with theta plasmid (trial 1).

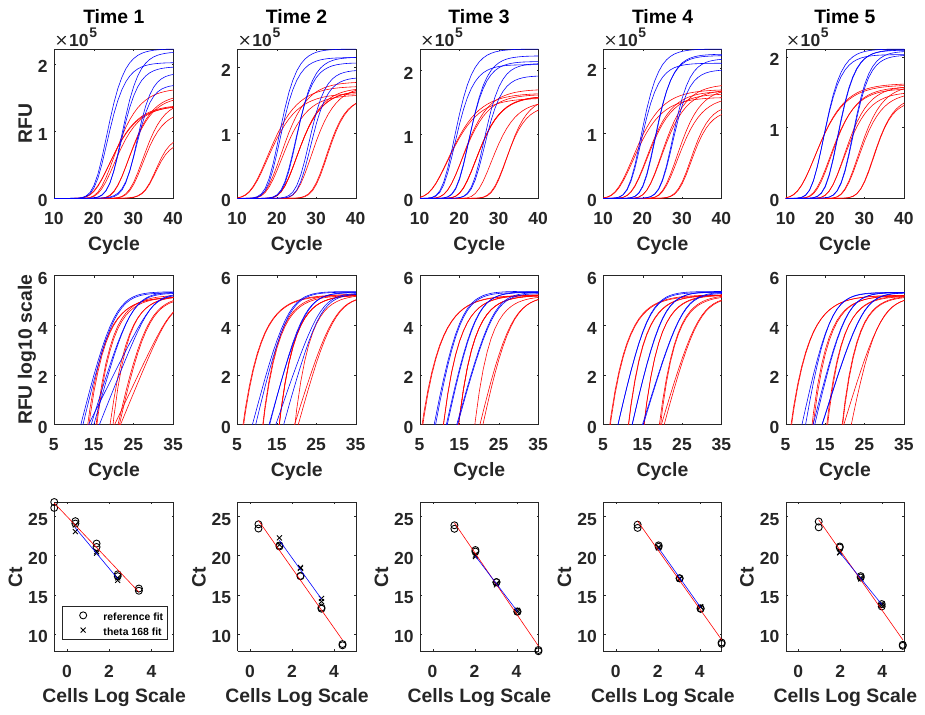

Supplement figure 7 qPCR data from B. subtilis cells with theta plasmid (trial 2).

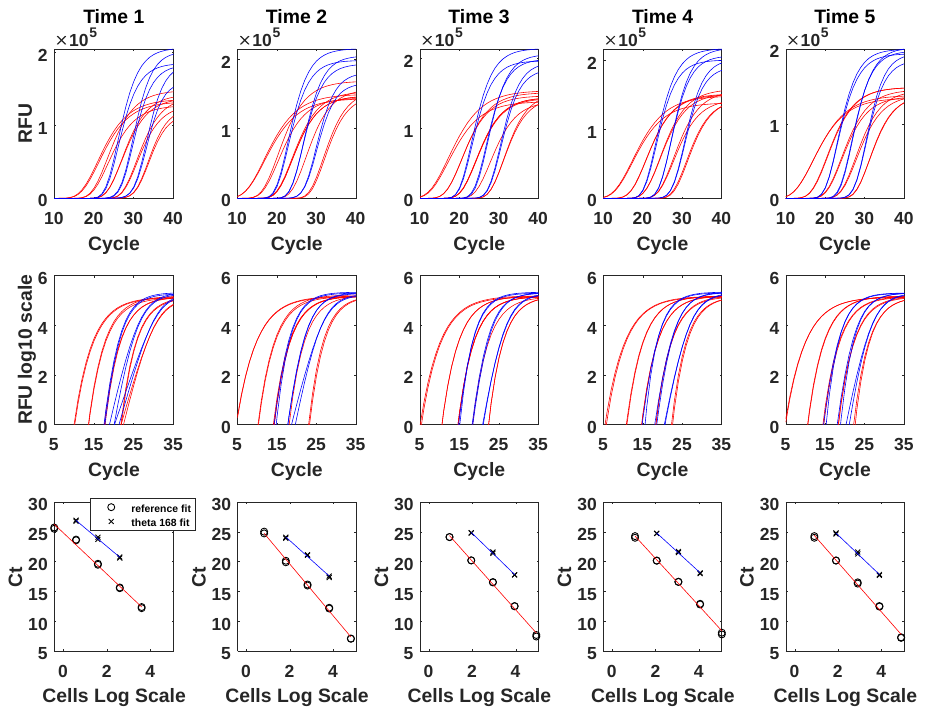

Supplement figure 8 qPCR data from B. subtilis cells with integrating plasmid (trial 1).

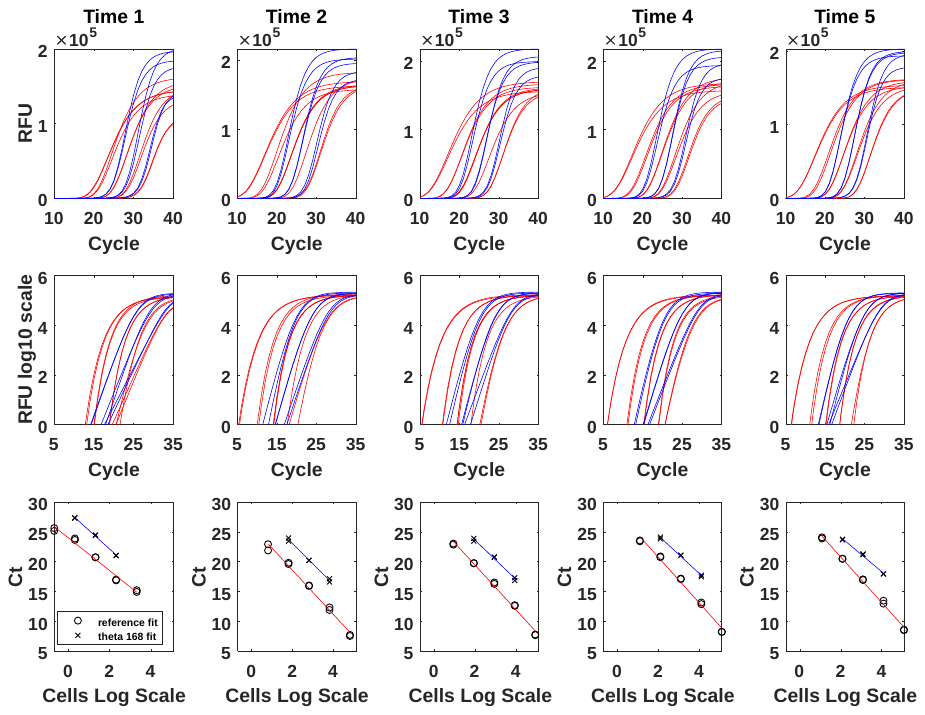

Supplement figure 9 qPCR data from B. subtilis cells with integrating plasmid (trial 2).

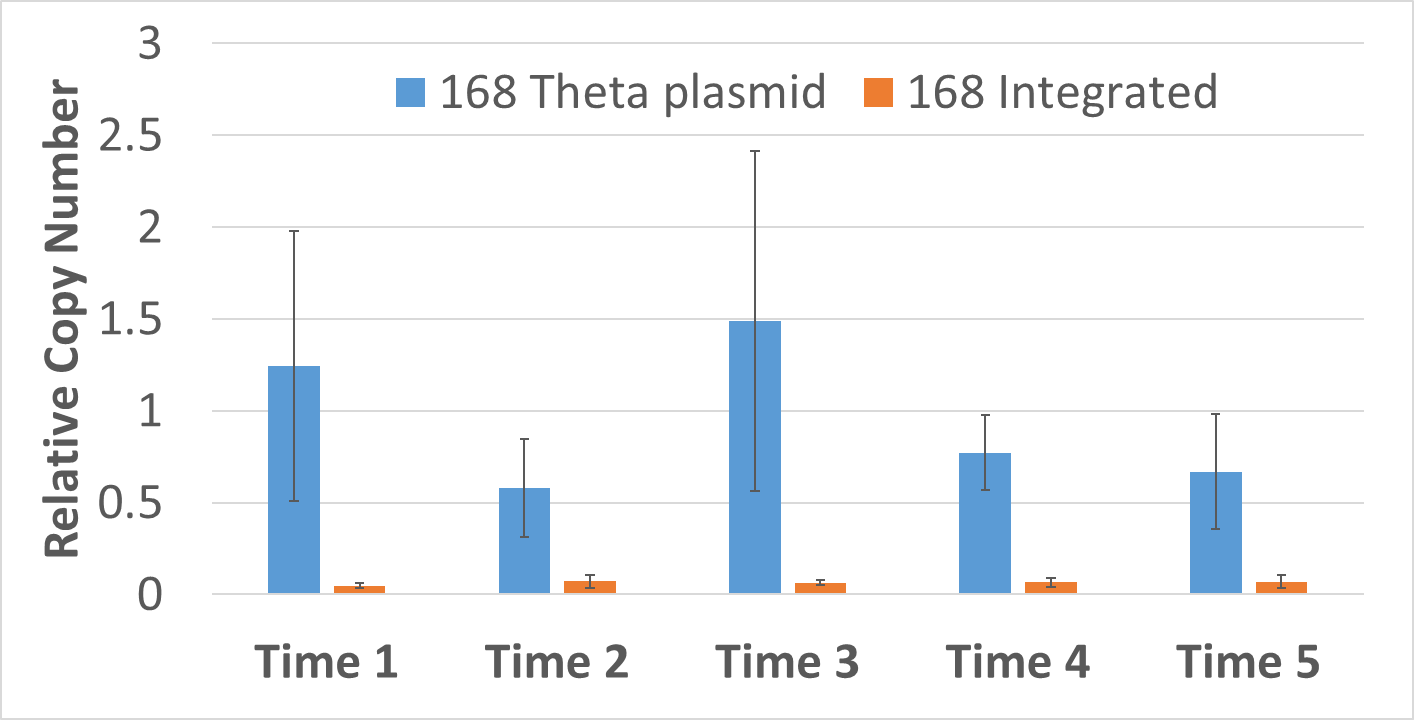

Supplement figure 10 Average relative copy number values of wild type cells carrying either a theta plasmid or integrated DNA harvested at different growth phases.

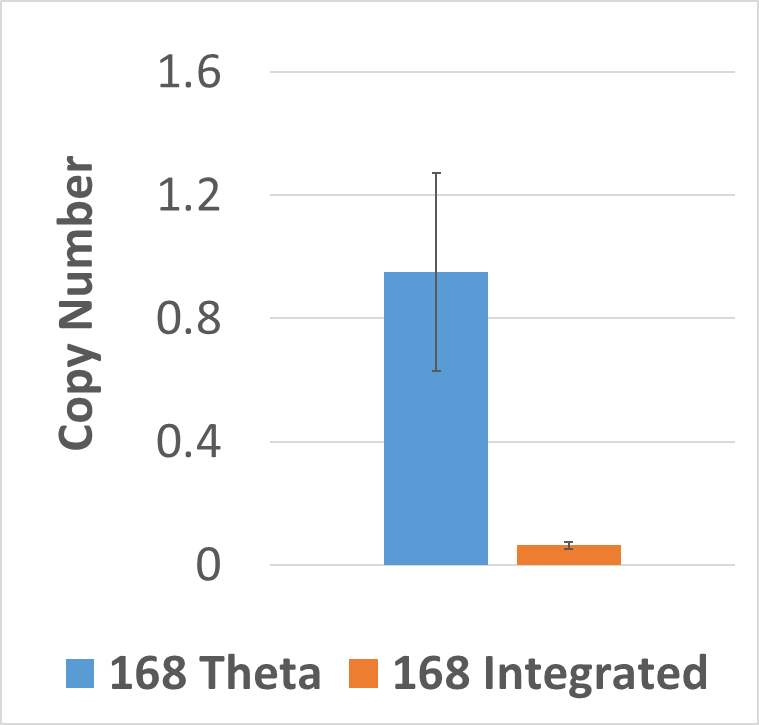

Supplement figure 11 Average, across all growth phases, relative copy number values of wild type cells carrying either a theta plasmid or integrated DNA harvested at different times during the growth.

Supplement table 3 RNA concentration after RNA purification

|  | Theta | |  | Integrating | |  |  |
| --- | --- | --- | --- | --- | --- | --- | --- |
|  | Trial 1 | Trial 2 | AVG | Trial 1 | Trial 2 |  |  |
| Time 1 | 38.298 | 46.915 | 42.6065 | 24.065 | 34.194 | 29.1295 | ng/ul |
| Time 2 | 52.504 | 58.531 | 55.5175 | 31.84 | 36.602 | 34.221 | ng/ul |
| Time 3 | 688.631 | 786.852 | 737.7415 | 32.277 | 31.107 | 31.692 | ng/ul |
| Time 4 | 58.19 | 58.58 | 58.385 | 56.29 | 57.222 | 56.756 | ng/ul |
| Time 5 | 166.547 | 185.935 | 176.241 | 218.629 | 219.712 | 219.1705 | ng/ul |

Supplement table 4 RNA concentration in RT-qPCR reactions (dilution replicates)

|  | **Theta** | | | **Integrating** | | |  |
| --- | --- | --- | --- | --- | --- | --- | --- |
|  | 0.1X | 0.001X | 0.00001X | 0.1X | 0.001X | 0.00001X |  |
| Time 1 | 4.26065 | 0.042607 | 0.000426 | 2.91295 | 0.02913 | 0.000291 | ng/ul |
| Time 2 | 5.55175 | 0.055518 | 0.000555 | 3.4221 | 0.034221 | 0.000342 | ng/ul |
| Time 3 | 73.77415 | 0.737742 | 0.007377 | 3.1692 | 0.031692 | 0.000317 | ng/ul |
| Time 4 | 5.8385 | 0.058385 | 0.000584 | 5.6756 | 0.056756 | 0.000568 | ng/ul |
| Time 5 | 17.6241 | 0.176241 | 0.001762 | 21.91705 | 0.219171 | 0.002192 | ng/ul |

Supplement table 5 Theophylline RNA normalized by total RNA in reaction (figure 5A)

| Theophylline RNA (copies) / RNA total (ng) | | | |
| --- | --- | --- | --- |
| Theta | | Integrating | |
| AVG | STDEV | AVG | STDEV |
| 10838033 | 129778.04 | 1342857.6 | 44901.413 |
| 3691818.1 | 449778.73 | 346364.57 | 22150.31 |
| 805934.65 | 46094.536 | 128756.53 | 14564.752 |
| 1161384.5 | 238352.27 | 64019.88 | 745.32271 |
| 646788.43 | 73095.961 | 17712.588 | 3300.6171 |

Supplement table 6 Riboswitch RNA normalized by total RNA in RT-qPCR reaction and DNA copy number (figure 5B)

| Theta | | | Integrating | | |
| --- | --- | --- | --- | --- | --- |
| Average | Error High | Error Low | Average | Error High | Error Low |
| 1205297.3 | 2494822 | 788080.41 | 2581425.6 | 5026655 | 1698184 |
| 410566.96 | 942079.19 | 238599.81 | 665829.62 | 1334812 | 424186.5 |
| 89627.964 | 193809.06 | 55920.884 | 247513.51 | 519129.5 | 149403.1 |
| 129157.52 | 318394.79 | 67931.103 | 123067.82 | 234588.5 | 82785.43 |
| 71929.319 | 163750.39 | 42221.238 | 34049.572 | 76112.74 | 18855.94 |

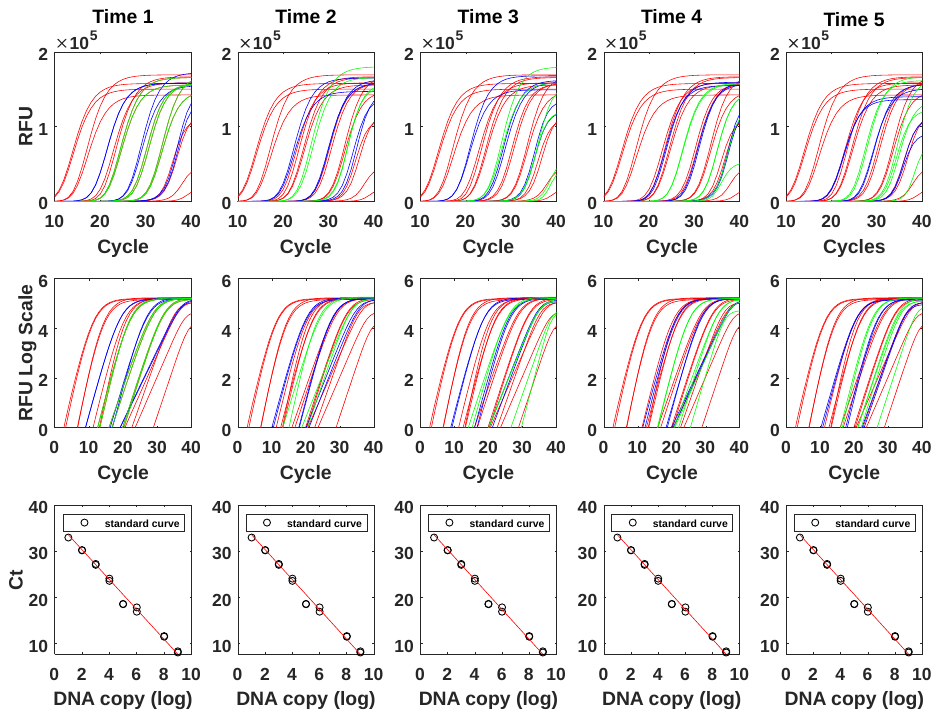

Supplement figure 12 workflow of RT-qPCR data processing

```
%coded on 11/15/2018 by DDT
%modified for experiment 6 on 01/24/2019 by DDT

%The code fluorescence data for all 4 channels from all wells and organizes it per cycle.
%Code requires raw data output from OneStep software in a separate .xlsx file
%Columns grouped in 4 per each well (B, G, Y, R). Order of wells dictated by the well matrix.

function qPCR_data_analyzer
clear
clc
tic
rRFU= readtable('Raw Data.xlsx');
rRFU = rRFU(8:7687,:); %trims the table to keep only numbers (no headings)

wellIDs  =rRFU(:,1);
cycleNum =rRFU(:,2);
Blue     =rRFU(:,3);
Green    =rRFU(:,4);
Yellow   =rRFU(:,5);
Red      =rRFU(:,6);

L=size(cycleNum); % total number of rows in the table
L=L(1,1);

tCycles=80; %total number of amplification cycles account for all types of cycles, denaturation/annealing/amplification

%create indexing arrays
wellMatrix={'A1';'A2';'A3';'A4';'A5';'A6';'A7';'A8';'A9';'A10';'A11';'A12';'B1';'B2';'B3';'B4';'B5';'B6';'B7';'B8';'B9';'B10';'B11';'B12';'C1';'C2';'C3';'C4';'C5';'C6';'C7';'C8';'C9';'C10';'C11';'C12';'D1';'D2';'D3';'D4';'D5';'D6';'D7';'D8';'D9';'D10';'D11';'D12';'E1';'E2';'E3';'E4';'E5';'E6';'E7';'E8';'E9';'E10';'E11';'E12';'F1';'F2';'F3';'F4';'F5';'F6';'F7';'F8';'F9';'F10';'F11';'F12';'G1';'G2';'G3';'G4';'G5';'G6';'G7';'G8';'G9';'G10';'G11';'G12';'H1';'H2';'H3';'H4';'H5';'H6';'H7';'H8';'H9';'H10';'H11';'H12'};
rawData=zeros(tCycles, 4, 96); %number of cycles | RFU for each color (4) | well id (96); will parse wells and cycles in a different function

for w=1:96 %untill all wells are assigned
c=1; %current row in the rawData matrix (to be filled). With max value = tCycles, total number of rows in the output matrix;
r=w; %r=current row number in matrix rRFU, with max=L. Indexing through raw data.
cName=cell2mat(wellMatrix(w)); %current well id that is being filled.
rawData(c,1:4,w)=cellfun(@str2num,table2cell(rRFU(w,3:6))); %manually fills the first row of the output table.
c=c+1; %moves to the next cycle since the first cycle has been filled manually in Line 33.
r=r+1; %moves to the next row in the raw data table since the first one has been used in Line 33
logical=zeros(L,1); %conditional statement to match well ids of <rRFU> and <wellMatrix>.

    for r=r:L %for all of the rows below this one
        logical(r,1)=strcmp(table2array(rRFU(r,1)),cName);
        if strcmp(table2array(rRFU(r,1)),cName)==1
           rawData(c,1:4,w)=cellfun(@str2num,table2cell(rRFU(r,3:6)));
           c=c+1;
        end
    end

end

%The data will be organized in 80 rows (one for each cycle)
%Four columns for each well, the order of wells is dictated by the well matrix.
csvwrite('Organized data.csv',rawData);
toc
end


%coded on 11/15/2018 by DDT
%V2 modified on 01/06/2018
%%Modification objective is to automatically fit every well and generate
%this matLab file must be located in the same folder as the data file
%______________________________________________________________________
%input data requirements: you need (2) files.
%qPCR data from AIB OneStepPlus exports data in the .xls file. Raw data
%from that file will be parsed and organized by the function
%"qPCR_data_analyzer.m". The output of that function is "data.csv" (see
%"qPCR_data_analyzer.m" for details on data structure in data.csv). After
%that, you will need legends that were assigned to this data in the
%OneStepPlus software. OneStepPlus takes data for all channels and all
%wells. To only keep necessary data this program will crossreference raw
%data from "data.csv" and the legened.
%(1) import data.csv using [~,~,<name>]=xlsread('data.csv');
%(2) import the legend from the original OneStepPlus output file using
%[~,~,<name>]=xlsread('<name of the OneStepPlus output file>', 'Sample
%Setup');
%system output is a structured array with columns: [well] [type] [dilution]
%[RFUprobe] [RFUrox]

%{
%EXAMPLES OF USAGE:
[~,~,data] = xlsread('organized data.csv'); %output from qPCR_data_analyzer
[~,~,legend] = xlsread('Experiment 4','Sample Setup');
qPCRdata=qPCR2mat(data, legend);
%}


function [qPCRdata]=qPCR2mat_V2(data, legend,Ct)
tic

%will need this at some point for crossreference
wellMatrix={'A1';'A2';'A3';'A4';'A5';'A6';'A7';'A8';'A9';'A10';'A11';'A12';'B1';'B2';'B3';'B4';'B5';'B6';'B7';'B8';'B9';'B10';'B11';'B12';'C1';'C2';'C3';'C4';'C5';'C6';'C7';'C8';'C9';'C10';'C11';'C12';'D1';'D2';'D3';'D4';'D5';'D6';'D7';'D8';'D9';'D10';'D11';'D12';'E1';'E2';'E3';'E4';'E5';'E6';'E7';'E8';'E9';'E10';'E11';'E12';'F1';'F2';'F3';'F4';'F5';'F6';'F7';'F8';'F9';'F10';'F11';'F12';'G1';'G2';'G3';'G4';'G5';'G6';'G7';'G8';'G9';'G10';'G11';'G12';'H1';'H2';'H3';'H4';'H5';'H6';'H7';'H8';'H9';'H10';'H11';'H12'};

%input RFU data
input = data;

%Parse the legend file to remove headers
vLength=length(legend);
N=find(strcmp(legend,'Well'));
legend=cell2table(legend(N+1:vLength,1:12));


%create a cell array of samples that contain: (1) name of the well
n=1;
for i=1:96

    if cellfun(@isempty,table2cell(legend(i,7)))==0
        allSamples(n,1)=table2cell(legend(i,1)); %name of the well
        %allSamples(n,2)=table2cell(legend(i,7)); %target name
        %allSamples(n,3)=table2cell(legend(i,12));%dilution
        n=n+1;

    end
end

%convert the reference cell array into a structured array
colHeadings={'well'};
allSamples=cell2struct(allSamples,colHeadings,2);


%organizing raw RFU data (BLUE and RED) into all 96 wells
%%change if you start using different dye or if the number of cycles is
%%different
rawData=zeros(80,2,96);
for i=1:96
    n=4*i-3; %every first column
    m=4*i;   %every fourth column

    rawData(1:80,1,i)=cell2mat(input(1:80,n)); %blue
    rawData(1:80,2,i)=cell2mat(input(1:80,m)); %red

end

%trim data
i=1:2:80; %every odd step (all denaturation steps removed)
rawData(i,:,:)=[];

%add RFU data to the structure
for i=1:length(allSamples)
    A=find(strcmp(wellMatrix,allSamples(i).well)); %row id
    allSamples(i).RFUprobe=rawData(:,1,A);
    allSamples(i).RFUrox=rawData(:,2,A);
end


%Loop through all data and attempt fitting it with 5pl
for i=1:length(allSamples)
   A=allSamples(i).RFUprobe(1);
   B=allSamples(i).RFUprobe(40);
   X=transpose(1:40);

    if A>0 && B>0 && B>A %if there is signal from the well (~negative RFU and final RFU > initial RFU)
       y=allSamples(i).RFUprobe;
       [xData, yData] = prepareCurveData( X, y );

       %customized guess parameters for upper and lower bounds:

       cMin=y(1)-y(1)*0.05; % 5 percent below min
       cMax=y(1)+y(1)*0.05; % 5 percent above min

       dMin=y(40)-y(40)*0.05;
       dMax=y(40)+y(40)*0.05;

       try
       ft = fittype( 'c+(d-c)/(1+exp(b*(x-e)))^f', 'independent', 'x', 'dependent', 'y' );
       opts = fitoptions( 'Method', 'NonlinearLeastSquares' );
       opts.Display = 'Off';
       opts.Lower = [-1 cMin dMin -Inf 0];
       %opts.StartPoint = [0.816675835667726 0.64689910993646 0.913375856139019 0.63235924622541 0.0975404049994095];
       opts.StartPoint = [0.816675835667726 0.8 0.913375856139019 0.63235924622541 0.0975404049994095];
       opts.Upper = [0 cMax dMax 40 100];
       [p, gof] = fit( xData, yData, ft, opts );
       catch
       p.b = 0; p.c = 0; p.d = 0; p.e = 0; p.f = 0; gof.rsquare = 0;
       end

       fitData=p.c+(p.d-p.c)./(1+exp(p.b.*(X-p.e))).^p.f;

       %fill in the structure
       allSamples(i).fitData=fitData;
       allSamples(i).rsqr=gof.rsquare;

       %Normalize by subtracting the first value
       [A]=allSamples(i).fitData;
       B=A-A(1);
       allSamples(i).Normalized=B; %add to the structure

        %convert to the log scale
        B=log10(B);
        allSamples(i).LogScale=B;

        %identify Ct for a given threshold by interpolation
        RFU= @(z) (p.c+(p.d-p.c)./(1+exp(p.b.*(z-p.e))).^p.f)-A(1); % eqn format, baseline normalized
        Zvec = linspace(1,40,1000); %defines independent var range
        RFUlog=log10(RFU(Zvec)); %initiates Y values Zvec range in the log10 format


        RFUlog=RFUlog(RFUlog > 0);  %adjusts the vector length to only positive values.
        startVal=1000-length(RFUlog)+1;

        bingo=spline(RFUlog',Zvec(startVal:1000)',Ct); %ids the Z value of that meets the Ct

        allSamples(i).Ct=bingo;
    end
end

%output is a single structure with substructures
qPCRdata.allSamples=allSamples;

toc %clock off
end


%This function plots the data
function [masterData]=plotqPCR
clc
clearvars
close all
tic
%load qPCRdata
[~,~,data] = xlsread('Organized data.csv');
[~,~,ss] = xlsread('Experiment 10_data','Sample Setup');
qPCRdata=qPCR2mat_V2(data, ss, 2.0);
qPCRdataT=qPCR2mat_V2(data, ss, 3.0);
%save('qPCRdata');

%%
%organize the data
REF=table('Size', [50,3],'VariableTypes', {'cell' 'cell' 'double'});
TH=table('Size', [30,3],'VariableTypes', {'cell' 'cell' 'double'});
%reference time 1
REF(1:2,1)={qPCRdata.allSamples(1).Normalized;qPCRdata.allSamples(2).Normalized}; %1X
REF(1:2,2)={qPCRdata.allSamples(1).LogScale; qPCRdata.allSamples(2).LogScale;};
REF(1:2,3)={qPCRdata.allSamples(1).Ct; qPCRdata.allSamples(2).Ct};

REF(3:4,1)={qPCRdata.allSamples(11).Normalized;qPCRdata.allSamples(12).Normalized}; %0.1X
REF(3:4,2)={qPCRdata.allSamples(11).LogScale; qPCRdata.allSamples(12).LogScale;};
REF(3:4,3)={qPCRdata.allSamples(11).Ct; qPCRdata.allSamples(12).Ct};

REF(5:6,1)={qPCRdata.allSamples(21).Normalized;qPCRdata.allSamples(22).Normalized}; %0.01X
REF(5:6,2)={qPCRdata.allSamples(21).LogScale; qPCRdata.allSamples(22).LogScale;};
REF(5:6,3)={qPCRdata.allSamples(21).Ct; qPCRdata.allSamples(22).Ct};

REF(7:8,1)={qPCRdata.allSamples(31).Normalized;qPCRdata.allSamples(32).Normalized}; %0.001X
REF(7:8,2)={qPCRdata.allSamples(31).LogScale; qPCRdata.allSamples(32).LogScale;};
REF(7:8,3)={qPCRdata.allSamples(31).Ct; qPCRdata.allSamples(32).Ct};

REF(9:10,1)={qPCRdata.allSamples(41).Normalized;qPCRdata.allSamples(42).Normalized}; %0.0001X
REF(9:10,2)={qPCRdata.allSamples(41).LogScale; qPCRdata.allSamples(42).LogScale;};
REF(9:10,3)={qPCRdata.allSamples(41).Ct; qPCRdata.allSamples(42).Ct};


%reference time 2
REF(11:12,1)={qPCRdata.allSamples(3).Normalized;qPCRdata.allSamples(4).Normalized}; %1X
REF(11:12,2)={qPCRdata.allSamples(3).LogScale; qPCRdata.allSamples(4).LogScale;};
REF(11:12,3)={qPCRdata.allSamples(3).Ct; qPCRdata.allSamples(4).Ct};

REF(13:14,1)={qPCRdata.allSamples(13).Normalized;qPCRdata.allSamples(14).Normalized}; %0.1X
REF(13:14,2)={qPCRdata.allSamples(13).LogScale; qPCRdata.allSamples(14).LogScale;};
REF(13:14,3)={qPCRdata.allSamples(13).Ct; qPCRdata.allSamples(14).Ct};

REF(15:16,1)={qPCRdata.allSamples(23).Normalized;qPCRdata.allSamples(24).Normalized}; %0.01X
REF(15:16,2)={qPCRdata.allSamples(23).LogScale; qPCRdata.allSamples(24).LogScale;};
REF(15:16,3)={qPCRdata.allSamples(23).Ct; qPCRdata.allSamples(24).Ct};

REF(17:18,1)={qPCRdata.allSamples(33).Normalized;qPCRdata.allSamples(34).Normalized}; %0.001X
REF(17:18,2)={qPCRdata.allSamples(33).LogScale; qPCRdata.allSamples(34).LogScale;};
REF(17:18,3)={qPCRdata.allSamples(33).Ct; qPCRdata.allSamples(34).Ct};

REF(19:20,1)={qPCRdata.allSamples(43).Normalized;qPCRdata.allSamples(44).Normalized}; %0.0001X
REF(19:20,2)={qPCRdata.allSamples(43).LogScale; qPCRdata.allSamples(44).LogScale;};
REF(19:20,3)={qPCRdata.allSamples(43).Ct; qPCRdata.allSamples(44).Ct};

%reference time 3
REF(21:22,1)={qPCRdata.allSamples(5).Normalized;qPCRdata.allSamples(6).Normalized}; %1X
REF(21:22,2)={qPCRdata.allSamples(5).LogScale; qPCRdata.allSamples(6).LogScale;};
REF(21:22,3)={qPCRdata.allSamples(5).Ct; qPCRdata.allSamples(6).Ct};

REF(23:24,1)={qPCRdata.allSamples(15).Normalized;qPCRdata.allSamples(16).Normalized}; %0.1X
REF(23:24,2)={qPCRdata.allSamples(15).LogScale; qPCRdata.allSamples(16).LogScale;};
REF(23:24,3)={qPCRdata.allSamples(15).Ct; qPCRdata.allSamples(16).Ct};

REF(25:26,1)={qPCRdata.allSamples(25).Normalized;qPCRdata.allSamples(26).Normalized}; %0.01X
REF(25:26,2)={qPCRdata.allSamples(25).LogScale; qPCRdata.allSamples(26).LogScale;};
REF(25:26,3)={qPCRdata.allSamples(25).Ct; qPCRdata.allSamples(26).Ct};

REF(27:28,1)={qPCRdata.allSamples(35).Normalized;qPCRdata.allSamples(36).Normalized}; %0.001X
REF(27:28,2)={qPCRdata.allSamples(35).LogScale; qPCRdata.allSamples(36).LogScale;};
REF(27:28,3)={qPCRdata.allSamples(35).Ct; qPCRdata.allSamples(36).Ct};

REF(29:30,1)={qPCRdata.allSamples(45).Normalized;qPCRdata.allSamples(46).Normalized}; %0.0001X
REF(29:30,2)={qPCRdata.allSamples(45).LogScale; qPCRdata.allSamples(46).LogScale;};
REF(29:30,3)={qPCRdata.allSamples(45).Ct; qPCRdata.allSamples(46).Ct};

%reference time 4
REF(31:32,1)={qPCRdata.allSamples(7).Normalized;qPCRdata.allSamples(8).Normalized}; %1X
REF(31:32,2)={qPCRdata.allSamples(7).LogScale; qPCRdata.allSamples(8).LogScale;};
REF(31:32,3)={qPCRdata.allSamples(7).Ct; qPCRdata.allSamples(8).Ct};

REF(33:34,1)={qPCRdata.allSamples(17).Normalized;qPCRdata.allSamples(18).Normalized}; %0.1X
REF(33:34,2)={qPCRdata.allSamples(17).LogScale; qPCRdata.allSamples(18).LogScale;};
REF(33:34,3)={qPCRdata.allSamples(17).Ct; qPCRdata.allSamples(18).Ct};

REF(35:36,1)={qPCRdata.allSamples(27).Normalized;qPCRdata.allSamples(28).Normalized}; %0.01X
REF(35:36,2)={qPCRdata.allSamples(27).LogScale; qPCRdata.allSamples(28).LogScale;};
REF(35:36,3)={qPCRdata.allSamples(27).Ct; qPCRdata.allSamples(28).Ct};

REF(37:38,1)={qPCRdata.allSamples(37).Normalized;qPCRdata.allSamples(38).Normalized}; %0.001X
REF(37:38,2)={qPCRdata.allSamples(37).LogScale; qPCRdata.allSamples(38).LogScale;};
REF(37:38,3)={qPCRdata.allSamples(37).Ct; qPCRdata.allSamples(38).Ct};

REF(39:40,1)={qPCRdata.allSamples(47).Normalized;qPCRdata.allSamples(48).Normalized}; %0.0001X
REF(39:40,2)={qPCRdata.allSamples(47).LogScale; qPCRdata.allSamples(48).LogScale;};
REF(39:40,3)={qPCRdata.allSamples(47).Ct; qPCRdata.allSamples(48).Ct};

%referrence time 5
REF(41:42,1)={qPCRdata.allSamples(9).Normalized;qPCRdata.allSamples(10).Normalized}; %1X
REF(41:42,2)={qPCRdata.allSamples(9).LogScale; qPCRdata.allSamples(10).LogScale;};
REF(41:42,3)={qPCRdata.allSamples(9).Ct; qPCRdata.allSamples(10).Ct};

REF(43:44,1)={qPCRdata.allSamples(19).Normalized;qPCRdata.allSamples(20).Normalized}; %0.1X
REF(43:44,2)={qPCRdata.allSamples(19).LogScale; qPCRdata.allSamples(20).LogScale;};
REF(43:44,3)={qPCRdata.allSamples(19).Ct; qPCRdata.allSamples(20).Ct};

REF(45:46,1)={qPCRdata.allSamples(29).Normalized;qPCRdata.allSamples(30).Normalized}; %0.01X
REF(45:46,2)={qPCRdata.allSamples(29).LogScale; qPCRdata.allSamples(30).LogScale;};
REF(45:46,3)={qPCRdata.allSamples(29).Ct; qPCRdata.allSamples(30).Ct};

REF(47:48,1)={qPCRdata.allSamples(39).Normalized;qPCRdata.allSamples(40).Normalized}; %0.001X
REF(47:48,2)={qPCRdata.allSamples(39).LogScale; qPCRdata.allSamples(40).LogScale;};
REF(47:48,3)={qPCRdata.allSamples(39).Ct; qPCRdata.allSamples(40).Ct};

REF(49:50,1)={qPCRdata.allSamples(49).Normalized;qPCRdata.allSamples(50).Normalized}; %0.0001X
REF(49:50,2)={qPCRdata.allSamples(49).LogScale; qPCRdata.allSamples(50).LogScale;};
REF(49:50,3)={qPCRdata.allSamples(49).Ct; qPCRdata.allSamples(50).Ct};

%%
%Theta time 1
TH(1:2,1)={qPCRdataT.allSamples(51).Normalized;qPCRdataT.allSamples(52).Normalized}; %0.1X
TH(1:2,2)={qPCRdataT.allSamples(51).LogScale; qPCRdataT.allSamples(52).LogScale;};
TH(1:2,3)={qPCRdataT.allSamples(51).Ct; qPCRdataT.allSamples(52).Ct};

TH(3:4,1)={qPCRdataT.allSamples(61).Normalized;qPCRdataT.allSamples(62).Normalized}; %0.01X
TH(3:4,2)={qPCRdataT.allSamples(61).LogScale; qPCRdataT.allSamples(62).LogScale;};
TH(3:4,3)={qPCRdataT.allSamples(61).Ct; qPCRdataT.allSamples(62).Ct};

TH(5:6,1)={qPCRdataT.allSamples(71).Normalized;qPCRdataT.allSamples(72).Normalized}; %0.001X
TH(5:6,2)={qPCRdataT.allSamples(71).LogScale; qPCRdataT.allSamples(72).LogScale;};
TH(5:6,3)={qPCRdataT.allSamples(71).Ct; qPCRdataT.allSamples(72).Ct};

%Theta time 2
TH(7:8,1)={qPCRdataT.allSamples(53).Normalized;qPCRdataT.allSamples(54).Normalized}; %0.1X
TH(7:8,2)={qPCRdataT.allSamples(53).LogScale; qPCRdataT.allSamples(54).LogScale;};
TH(7:8,3)={qPCRdataT.allSamples(53).Ct; qPCRdataT.allSamples(54).Ct};

TH(9:10,1)={qPCRdataT.allSamples(63).Normalized;qPCRdataT.allSamples(64).Normalized}; %0.01X
TH(9:10,2)={qPCRdataT.allSamples(63).LogScale; qPCRdataT.allSamples(64).LogScale;};
TH(9:10,3)={qPCRdataT.allSamples(63).Ct; qPCRdataT.allSamples(64).Ct};

TH(11:12,1)={qPCRdataT.allSamples(73).Normalized;qPCRdataT.allSamples(74).Normalized}; %0.001X
TH(11:12,2)={qPCRdataT.allSamples(73).LogScale; qPCRdataT.allSamples(74).LogScale;};
TH(11:12,3)={qPCRdataT.allSamples(73).Ct; qPCRdataT.allSamples(74).Ct};

%Theta time 3
TH(13:14,1)={qPCRdataT.allSamples(55).Normalized;qPCRdataT.allSamples(56).Normalized}; %0.1X
TH(13:14,2)={qPCRdataT.allSamples(55).LogScale; qPCRdataT.allSamples(56).LogScale;};
TH(13:14,3)={qPCRdataT.allSamples(55).Ct; qPCRdataT.allSamples(56).Ct};

TH(15:16,1)={qPCRdataT.allSamples(65).Normalized;qPCRdataT.allSamples(66).Normalized}; %0.01X
TH(15:16,2)={qPCRdataT.allSamples(65).LogScale; qPCRdataT.allSamples(66).LogScale;};
TH(15:16,3)={qPCRdataT.allSamples(65).Ct; qPCRdataT.allSamples(66).Ct};

TH(17:18,1)={qPCRdataT.allSamples(75).Normalized;qPCRdataT.allSamples(76).Normalized}; %0.001X
TH(17:18,2)={qPCRdataT.allSamples(75).LogScale; qPCRdataT.allSamples(76).LogScale;};
TH(17:18,3)={qPCRdataT.allSamples(75).Ct; qPCRdataT.allSamples(76).Ct};

%Theta time 4
TH(19:20,1)={qPCRdataT.allSamples(57).Normalized;qPCRdataT.allSamples(58).Normalized}; %0.1X
TH(19:20,2)={qPCRdataT.allSamples(57).LogScale; qPCRdataT.allSamples(58).LogScale;};
TH(19:20,3)={qPCRdataT.allSamples(57).Ct; qPCRdataT.allSamples(58).Ct};

TH(21:22,1)={qPCRdataT.allSamples(67).Normalized;qPCRdataT.allSamples(68).Normalized}; %0.01X
TH(21:22,2)={qPCRdataT.allSamples(67).LogScale; qPCRdataT.allSamples(68).LogScale;};
TH(21:22,3)={qPCRdataT.allSamples(67).Ct; qPCRdataT.allSamples(68).Ct};

TH(23:24,1)={qPCRdataT.allSamples(77).Normalized;qPCRdataT.allSamples(78).Normalized}; %0.001X
TH(23:24,2)={qPCRdataT.allSamples(77).LogScale; qPCRdataT.allSamples(78).LogScale;};
TH(23:24,3)={qPCRdataT.allSamples(77).Ct; qPCRdataT.allSamples(78).Ct};

%Theta time 5
TH(25:26,1)={qPCRdataT.allSamples(59).Normalized;qPCRdataT.allSamples(60).Normalized}; %0.1X
TH(25:26,2)={qPCRdataT.allSamples(59).LogScale; qPCRdataT.allSamples(60).LogScale;};
TH(25:26,3)={qPCRdataT.allSamples(59).Ct; qPCRdataT.allSamples(60).Ct};

TH(27:28,1)={qPCRdataT.allSamples(69).Normalized;qPCRdataT.allSamples(70).Normalized}; %0.01X
TH(27:28,2)={qPCRdataT.allSamples(69).LogScale; qPCRdataT.allSamples(70).LogScale;};
TH(27:28,3)={qPCRdataT.allSamples(69).Ct; qPCRdataT.allSamples(70).Ct};

TH(29:30,1)={qPCRdataT.allSamples(79).Normalized;qPCRdataT.allSamples(80).Normalized}; %0.001X
TH(29:30,2)={qPCRdataT.allSamples(79).LogScale; qPCRdataT.allSamples(80).LogScale;};
TH(29:30,3)={qPCRdataT.allSamples(79).Ct; qPCRdataT.allSamples(80).Ct};
%%
%Concentrations
CELL1=[2059.125; 205.9125; 20.59125; 2.059125; 0.2059125];

CELL2=[63547.5; 6354.75; 635.475; 63.5475; 6.35475];

CELL3=[87285; 8728.5; 872.85; 87.285; 8.7285];

CELL4=[125117.5; 12511.75; 1251.175; 125.1175; 12.51175];

CELL5=[120443.1; 12044.31; 1204.431; 120.4431; 12.04431];

%%
%create a matrix for fit values
masterData=zeros(5,6);
% (1) slope of the ref fit
% (2) y-int of the ref fit
% (3) rsqfor the ref fit
% (4-6) unknown fit
%These parameters will be recorded for each time point

%%
%Plot all figures
figure
%Time 1
ax1=subplot(3,5,11); %Standard curve
Xvec(1:5)=log10(CELL1);
Xvec(6:10)=Xvec;

Yvec([1 6])=table2array(REF(1:2,3));
Yvec([2 7])=table2array(REF(3:4,3));
Yvec([3 8])=table2array(REF(5:6,3));
Yvec([4 9])=table2array(REF(7:8,3));
Yvec([5 10])=table2array(REF(9:10,3));

XvecT(1:3)=log10(CELL1(2:4));
XvecT(4:6)=XvecT;

YvecT([1 4])=table2array(TH(1:2,3));
YvecT([2 5])=table2array(TH(3:4,3));
YvecT([3 6])=table2array(TH(5:6,3));

plot(Xvec,Yvec,'o', 'Color', 'black');
hold on
plot(XvecT,YvecT,'X', 'Color', 'black');
hold on

%linear fit (plot)
[xData, yData] = prepareCurveData( Xvec, Yvec);
ft = fittype( 'm*x+b', 'independent', 'x', 'dependent', 'y' );
[p, gof] = fit( xData, yData, ft);
YvecPlot=p.m*Xvec+p.b;
plot(Xvec,YvecPlot, 'Color', 'red');

%import the fit parameters into the matrix

masterData(1,1)=p.m;
masterData(1,2)=p.b;
masterData(1,3)=gof.rsquare;

hold on
[xData, yData] = prepareCurveData( XvecT, YvecT);
ft = fittype( 'm*x+b', 'independent', 'x', 'dependent', 'y' );
[p, gof] = fit( xData, yData, ft);
YvecPlot=p.m*XvecT+p.b;
plot(XvecT,YvecPlot, 'Color', 'blue');
legend('reference fit', 'theta 168 fit');
xlabel('[Cells (log10)]');
ylabel('Ct');

masterData(1,4)=p.m;
masterData(1,5)=p.b;
masterData(1,6)=gof.rsquare;

%%
subplot(3,5,6) %This is the log data;
Xvec=transpose(1:40);

%plot all of the reference curves
plot(Xvec,cell2mat(REF{1,2}),Xvec,cell2mat(REF{2,2}), 'Color', 'r'); %1X REF
hold on
plot(Xvec,cell2mat(REF{3,2}),Xvec,cell2mat(REF{4,2}), 'Color', 'r'); %0.1X REF
hold on
plot(Xvec,cell2mat(REF{5,2}),Xvec,cell2mat(REF{6,2}), 'Color', 'r'); %0.01X REF
hold on
plot(Xvec,cell2mat(REF{7,2}),Xvec,cell2mat(REF{8,2}), 'Color', 'r'); %0.001X REF
hold on
plot(Xvec,cell2mat(REF{9,2}),Xvec,cell2mat(REF{10,2}), 'Color', 'r'); %0.0001X REF
hold on

%plot all of the theta curves
plot(Xvec,cell2mat(TH{1,2}),Xvec,cell2mat(TH{2,2}), 'Color', 'b'); %0.1X REF
hold on
plot(Xvec,cell2mat(TH{3,2}),Xvec,cell2mat(TH{4,2}), 'Color', 'b'); %0.01X REF
hold on
plot(Xvec,cell2mat(TH{5,2}),Xvec,cell2mat(TH{6,2}), 'Color', 'b'); %0.001X REF
hold on

%adjust plot properties
ylim([0 6]);
xlim([5 35])
xticks(0:5:40);
xlabel('Cycle');
ylabel('RFU log10 scale');

%%

subplot(3,5,1) %This is the linear amplification data
Xvec=transpose(1:40);

%plot all of the reference curves
plot(Xvec,cell2mat(REF{1,1}),Xvec,cell2mat(REF{2,1}), 'Color', 'r'); %1X REF
hold on
plot(Xvec,cell2mat(REF{3,1}),Xvec,cell2mat(REF{4,1}), 'Color', 'r'); %0.1X REF
hold on
plot(Xvec,cell2mat(REF{5,1}),Xvec,cell2mat(REF{6,1}), 'Color', 'r'); %0.01X REF
hold on
plot(Xvec,cell2mat(REF{7,1}),Xvec,cell2mat(REF{8,1}), 'Color', 'r'); %0.001X REF
hold on
plot(Xvec,cell2mat(REF{9,1}),Xvec,cell2mat(REF{10,1}), 'Color', 'r'); %0.001X REF
hold on

%plot all of the theta curves
plot(Xvec,cell2mat(TH{1,1}),Xvec,cell2mat(TH{2,1}), 'Color', 'b'); %0.1X REF
hold on
plot(Xvec,cell2mat(TH{3,1}),Xvec,cell2mat(TH{4,1}), 'Color', 'b'); %0.01X REF
hold on
plot(Xvec,cell2mat(TH{5,1}),Xvec,cell2mat(TH{6,1}), 'Color', 'b'); %0.001X REF
hold on


%adjust plot properties
xlim([10 40])
xticks(10:5:40);
xlabel('Cycle');
ylabel('RFU');
title('Time 1');

%%
%Time 2
ax2=subplot(3,5,12); %Standard curve
Xvec=[];
XvecT=[];
Xvec(1:5)=log10(CELL2);
Xvec(6:10)=Xvec;

Yvec([1 6])=table2array(REF(11:12,3));
Yvec([2 7])=table2array(REF(13:14,3));
Yvec([3 8])=table2array(REF(15:16,3));
Yvec([4 9])=table2array(REF(17:18,3));
Yvec([5 10])=table2array(REF(19:20,3));

XvecT(1:3)=log10(CELL2(2:4));
XvecT(4:6)=XvecT;

YvecT([1 4])=table2array(TH(7:8,3));
YvecT([2 5])=table2array(TH(9:10,3));
YvecT([3 6])=table2array(TH(11:12,3));

plot(Xvec,Yvec,'o', 'Color', 'black');
hold on
plot(XvecT,YvecT,'X', 'Color', 'black');
hold on

%linear fit (plot)
[xData, yData] = prepareCurveData( Xvec, Yvec);
ft = fittype( 'm*x+b', 'independent', 'x', 'dependent', 'y' );
[p, gof] = fit( xData, yData, ft);
YvecPlot=p.m*Xvec+p.b;
plot(Xvec,YvecPlot, 'Color', 'red');

%import the fit parameters into the matrix

masterData(2,1)=p.m;
masterData(2,2)=p.b;
masterData(2,3)=gof.rsquare;

hold on
[xData, yData] = prepareCurveData( XvecT, YvecT);
ft = fittype( 'm*x+b', 'independent', 'x', 'dependent', 'y' );
[p, gof] = fit( xData, yData, ft);
YvecPlot=p.m*XvecT+p.b;
plot(XvecT,YvecPlot, 'Color', 'blue');
legend('reference fit', 'theta 168 fit');
xlabel('[Cells (log10)]');
ylabel('Ct');

masterData(2,4)=p.m;
masterData(2,5)=p.b;
masterData(2,6)=gof.rsquare;

%%
subplot(3,5,7) %This is the log data;
Xvec=transpose(1:40);

%plot all of the reference curves
plot(Xvec,cell2mat(REF{11,2}),Xvec,cell2mat(REF{12,2}), 'Color', 'r'); %1X REF
hold on
plot(Xvec,cell2mat(REF{13,2}),Xvec,cell2mat(REF{14,2}), 'Color', 'r'); %0.1X REF
hold on
plot(Xvec,cell2mat(REF{15,2}),Xvec,cell2mat(REF{16,2}), 'Color', 'r'); %0.01X REF
hold on
plot(Xvec,cell2mat(REF{16,2}),Xvec,cell2mat(REF{18,2}), 'Color', 'r'); %0.001X REF
hold on
plot(Xvec,cell2mat(REF{19,2}),Xvec,cell2mat(REF{20,2}), 'Color', 'r'); %0.0001X REF
hold on

%plot all of the theta curves
plot(Xvec,cell2mat(TH{7,2}),Xvec,cell2mat(TH{8,2}), 'Color', 'b'); %1X REF
hold on
plot(Xvec,cell2mat(TH{9,2}),Xvec,cell2mat(TH{10,2}), 'Color', 'b'); %0.1X REF
hold on
plot(Xvec,cell2mat(TH{11,2}),Xvec,cell2mat(TH{12,2}), 'Color', 'b'); %0.01X REF
hold on


%adjust plot properties
ylim([0 6]);
xlim([5 35])
xticks(0:5:40);
xlabel('Cycle');

%%

subplot(3,5,2) %This is the linear amplification data
Xvec=transpose(1:40);

%plot all of the reference curves
plot(Xvec,cell2mat(REF{11,1}),Xvec,cell2mat(REF{12,1}), 'Color', 'r'); %1X REF
hold on
plot(Xvec,cell2mat(REF{13,1}),Xvec,cell2mat(REF{14,1}), 'Color', 'r'); %0.1X REF
hold on
plot(Xvec,cell2mat(REF{15,1}),Xvec,cell2mat(REF{16,1}), 'Color', 'r'); %0.01X REF
hold on
plot(Xvec,cell2mat(REF{16,1}),Xvec,cell2mat(REF{18,1}), 'Color', 'r'); %0.001X REF
hold on
plot(Xvec,cell2mat(REF{19,1}),Xvec,cell2mat(REF{20,1}), 'Color', 'r'); %0.0001X REF
hold on

%plot all of the theta curves
plot(Xvec,cell2mat(TH{7,1}),Xvec,cell2mat(TH{8,1}), 'Color', 'b'); %1X REF
hold on
plot(Xvec,cell2mat(TH{9,1}),Xvec,cell2mat(TH{10,1}), 'Color', 'b'); %0.1X REF
hold on
plot(Xvec,cell2mat(TH{11,1}),Xvec,cell2mat(TH{12,1}), 'Color', 'b'); %0.01X REF
hold on

%adjust plot properties
xlim([10 40])
xticks(10:5:40);
xlabel('Cycle');
title('Time 2');

%%
%Time 3
ax3=subplot(3,5,13); %Standard curve
Xvec=[];
XvecT=[];

Xvec(1:5)=log10(CELL3);
Xvec(6:10)=Xvec;

Yvec([1 6])=table2array(REF(21:22,3));
Yvec([2 7])=table2array(REF(23:24,3));
Yvec([3 8])=table2array(REF(25:26,3));
Yvec([4 9])=table2array(REF(27:28,3));
Yvec([5 10])=table2array(REF(29:30,3));

XvecT(1:3)=log10(CELL3(2:4));
XvecT(4:6)=XvecT;

YvecT([1 4])=table2array(TH(13:14,3));
YvecT([2 5])=table2array(TH(15:16,3));
YvecT([3 6])=table2array(TH(17:18,3));


plot(Xvec,Yvec,'o', 'Color', 'black');
hold on
plot(XvecT,YvecT,'X', 'Color', 'black');
hold on

%linear fit (plot)
[xData, yData] = prepareCurveData( Xvec, Yvec);
ft = fittype( 'm*x+b', 'independent', 'x', 'dependent', 'y' );
[p, gof] = fit( xData, yData, ft);
YvecPlot=p.m*Xvec+p.b;
plot(Xvec,YvecPlot, 'Color', 'red');

%import the fit parameters into the matrix

masterData(3,1)=p.m;
masterData(3,2)=p.b;
masterData(3,3)=gof.rsquare;

hold on
[xData, yData] = prepareCurveData( XvecT, YvecT);
ft = fittype( 'm*x+b', 'independent', 'x', 'dependent', 'y' );
[p, gof] = fit( xData, yData, ft);
YvecPlot=p.m*XvecT+p.b;
plot(XvecT,YvecPlot, 'Color', 'blue');
legend('reference fit', 'theta 168 fit');
xlabel('[Cells (log10)]');
ylabel('Ct');

masterData(3,4)=p.m;
masterData(3,5)=p.b;
masterData(3,6)=gof.rsquare;

%%
subplot(3,5,8) %This is the log data;
Xvec=transpose(1:40);

%plot all of the reference curves
plot(Xvec,cell2mat(REF{21,2}),Xvec,cell2mat(REF{22,2}), 'Color', 'r'); %1X REF
hold on
plot(Xvec,cell2mat(REF{23,2}),Xvec,cell2mat(REF{24,2}), 'Color', 'r'); %0.1X REF
hold on
plot(Xvec,cell2mat(REF{25,2}),Xvec,cell2mat(REF{26,2}), 'Color', 'r'); %0.01X REF
hold on
plot(Xvec,cell2mat(REF{27,2}),Xvec,cell2mat(REF{26,2}), 'Color', 'r'); %0.001X REF
hold on
plot(Xvec,cell2mat(REF{29,2}),Xvec,cell2mat(REF{30,2}), 'Color', 'r'); %0.001X REF
hold on

%plot all of the theta curves
plot(Xvec,cell2mat(TH{13,2}),Xvec,cell2mat(TH{14,2}), 'Color', 'b'); %1X REF
hold on
plot(Xvec,cell2mat(TH{15,2}),Xvec,cell2mat(TH{16,2}), 'Color', 'b'); %0.1X REF
hold on
plot(Xvec,cell2mat(TH{17,2}),Xvec,cell2mat(TH{18,2}), 'Color', 'b'); %0.01X REF
hold on

%adjust plot properties
ylim([0 6]);
xlim([5 35])
xticks(0:5:40);
xlabel('Cycle');

%%

subplot(3,5,3) %This is the linear amplification data
Xvec=transpose(1:40);

%plot all of the reference curves
plot(Xvec,cell2mat(REF{21,1}),Xvec,cell2mat(REF{22,1}), 'Color', 'r'); %1X REF
hold on
plot(Xvec,cell2mat(REF{23,1}),Xvec,cell2mat(REF{24,1}), 'Color', 'r'); %0.1X REF
hold on
plot(Xvec,cell2mat(REF{25,1}),Xvec,cell2mat(REF{26,1}), 'Color', 'r'); %0.01X REF
hold on
plot(Xvec,cell2mat(REF{27,1}),Xvec,cell2mat(REF{26,1}), 'Color', 'r'); %0.001X REF
hold on
plot(Xvec,cell2mat(REF{29,1}),Xvec,cell2mat(REF{30,1}), 'Color', 'r'); %0.001X REF
hold on

%plot all of the theta curves
plot(Xvec,cell2mat(TH{13,1}),Xvec,cell2mat(TH{14,1}), 'Color', 'b'); %1X REF
hold on
plot(Xvec,cell2mat(TH{15,1}),Xvec,cell2mat(TH{16,1}), 'Color', 'b'); %0.1X REF
hold on
plot(Xvec,cell2mat(TH{17,1}),Xvec,cell2mat(TH{18,1}), 'Color', 'b'); %0.01X REF
hold on

%adjust plot properties
xlim([10 40])
xticks(10:5:40);
xlabel('Cycle');
title('Time 3');

%%
%Time 4
ax4=subplot(3,5,14); %Standard curve
Xvec=[];
XvecT=[];

Xvec(1:5)=log10(CELL4);
Xvec(6:10)=Xvec;

Yvec([1 6])=table2array(REF(31:32,3));
Yvec([2 7])=table2array(REF(33:34,3));
Yvec([3 8])=table2array(REF(35:36,3));
Yvec([4 9])=table2array(REF(37:38,3));
Yvec([5 10])=table2array(REF(39:40,3));

XvecT(1:3)=log10(CELL4(2:4));
XvecT(4:6)=XvecT;

YvecT([1 4])=table2array(TH(19:20,3));
YvecT([2 5])=table2array(TH(21:22,3));
YvecT([3 6])=table2array(TH(23:24,3));


plot(Xvec,Yvec,'o', 'Color', 'black');
hold on
plot(XvecT,YvecT,'X', 'Color', 'black');
hold on

%linear fit (plot)
[xData, yData] = prepareCurveData( Xvec, Yvec);
ft = fittype( 'm*x+b', 'independent', 'x', 'dependent', 'y' );
[p, gof] = fit( xData, yData, ft);
YvecPlot=p.m*Xvec+p.b;
plot(Xvec,YvecPlot, 'Color', 'red');

%import the fit parameters into the matrix

masterData(4,1)=p.m;
masterData(4,2)=p.b;
masterData(4,3)=gof.rsquare;

hold on
[xData, yData] = prepareCurveData( XvecT, YvecT);
ft = fittype( 'm*x+b', 'independent', 'x', 'dependent', 'y' );
[p, gof] = fit( xData, yData, ft);
YvecPlot=p.m*XvecT+p.b;
plot(XvecT,YvecPlot, 'Color', 'blue');
legend('reference fit', 'theta 168 fit');
xlabel('[Cells (log10)]');
ylabel('Ct');

masterData(4,4)=p.m;
masterData(4,5)=p.b;
masterData(4,6)=gof.rsquare;

%%
subplot(3,5,9) %This is the log data;
Xvec=transpose(1:40);

%plot all of the reference curves
plot(Xvec,cell2mat(REF{31,2}),Xvec,cell2mat(REF{32,2}), 'Color', 'r'); %1X REF
hold on
plot(Xvec,cell2mat(REF{33,2}),Xvec,cell2mat(REF{34,2}), 'Color', 'r'); %0.1X REF
hold on
plot(Xvec,cell2mat(REF{35,2}),Xvec,cell2mat(REF{36,2}), 'Color', 'r'); %0.01X REF
hold on
plot(Xvec,cell2mat(REF{37,2}),Xvec,cell2mat(REF{38,2}), 'Color', 'r'); %0.001X REF
hold on
plot(Xvec,cell2mat(REF{39,2}),Xvec,cell2mat(REF{40,2}), 'Color', 'r'); %0.0001X REF
hold on

%plot all of the theta curves
plot(Xvec,cell2mat(TH{19,2}),Xvec,cell2mat(TH{20,2}), 'Color', 'b'); %0.1X REF
hold on
plot(Xvec,cell2mat(TH{21,2}),Xvec,cell2mat(TH{22,2}), 'Color', 'b'); %0.01X REF
hold on
plot(Xvec,cell2mat(TH{23,2}),Xvec,cell2mat(TH{24,2}), 'Color', 'b'); %0.001X REF
hold on


%adjust plot properties
ylim([0 6]);
xlim([5 35])
xticks(0:5:40);
xlabel('Cycle');


%%

subplot(3,5,4) %This is the linear amplification data
Xvec=transpose(1:40);

%plot all of the reference curves
plot(Xvec,cell2mat(REF{31,1}),Xvec,cell2mat(REF{32,1}), 'Color', 'r'); %1X REF
hold on
plot(Xvec,cell2mat(REF{33,1}),Xvec,cell2mat(REF{34,1}), 'Color', 'r'); %0.1X REF
hold on
plot(Xvec,cell2mat(REF{35,1}),Xvec,cell2mat(REF{36,1}), 'Color', 'r'); %0.01X REF
hold on
plot(Xvec,cell2mat(REF{37,1}),Xvec,cell2mat(REF{38,1}), 'Color', 'r'); %0.001X REF
hold on
plot(Xvec,cell2mat(REF{39,1}),Xvec,cell2mat(REF{40,1}), 'Color', 'r'); %0.0001X REF
hold on

%plot all of the theta curves
plot(Xvec,cell2mat(TH{19,1}),Xvec,cell2mat(TH{20,1}), 'Color', 'b'); %0.1X REF
hold on
plot(Xvec,cell2mat(TH{21,1}),Xvec,cell2mat(TH{22,1}), 'Color', 'b'); %0.01X REF
hold on
plot(Xvec,cell2mat(TH{23,1}),Xvec,cell2mat(TH{24,1}), 'Color', 'b'); %0.001X REF
hold on

%adjust plot properties
xlim([10 40])
xticks(10:5:40);

xlabel('Cycle');
title('Time 4');

%%
%Time 5
ax5=subplot(3,5,15); %Standard curve
Xvec=[];
XvecT=[];

Xvec(1:5)=log10(CELL5);
Xvec(6:10)=Xvec;

Yvec([1 6])=table2array(REF(41:42,3));
Yvec([2 7])=table2array(REF(43:44,3));
Yvec([3 8])=table2array(REF(45:46,3));
Yvec([4 9])=table2array(REF(47:48,3));
Yvec([5 10])=table2array(REF(49:50,3));

XvecT(1:3)=log10(CELL5(2:4));
XvecT(4:6)=XvecT;

YvecT([1 4])=table2array(TH(25:26,3));
YvecT([2 5])=table2array(TH(27:28,3));
YvecT([3 6])=table2array(TH(29:30,3));


plot(Xvec,Yvec,'o', 'Color', 'black');
hold on
plot(XvecT,YvecT,'X', 'Color', 'black');
hold on

%linear fit (plot)
[xData, yData] = prepareCurveData( Xvec, Yvec);
ft = fittype( 'm*x+b', 'independent', 'x', 'dependent', 'y' );
[p, gof] = fit( xData, yData, ft);
YvecPlot=p.m*Xvec+p.b;
plot(Xvec,YvecPlot, 'Color', 'red');

%import the fit parameters into the matrix

masterData(5,1)=p.m;
masterData(5,2)=p.b;
masterData(5,3)=gof.rsquare;

hold on
[xData, yData] = prepareCurveData( XvecT, YvecT);
ft = fittype( 'm*x+b', 'independent', 'x', 'dependent', 'y' );
[p, gof] = fit( xData, yData, ft);
YvecPlot=p.m*XvecT+p.b;
plot(XvecT,YvecPlot, 'Color', 'blue');
legend('reference fit', 'theta 168 fit');
xlabel('[Cells (log10)]');
ylabel('Ct');

masterData(5,4)=p.m;
masterData(5,5)=p.b;
masterData(5,6)=gof.rsquare;

%%
subplot(3,5,10) %This is the log data;
Xvec=transpose(1:40);

%plot all of the reference curves
plot(Xvec,cell2mat(REF{41,2}),Xvec,cell2mat(REF{42,2}), 'Color', 'r'); %1X REF
hold on
plot(Xvec,cell2mat(REF{43,2}),Xvec,cell2mat(REF{44,2}), 'Color', 'r'); %0.1X REF
hold on
plot(Xvec,cell2mat(REF{45,2}),Xvec,cell2mat(REF{46,2}), 'Color', 'r'); %0.01X REF
hold on
plot(Xvec,cell2mat(REF{47,2}),Xvec,cell2mat(REF{48,2}), 'Color', 'r'); %0.001X REF
hold on
plot(Xvec,cell2mat(REF{49,2}),Xvec,cell2mat(REF{50,2}), 'Color', 'r'); %0.001X REF
hold on

%plot all of the theta curves
plot(Xvec,cell2mat(TH{25,2}),Xvec,cell2mat(TH{26,2}), 'Color', 'b'); %1X REF
hold on
plot(Xvec,cell2mat(TH{27,2}),Xvec,cell2mat(TH{28,2}), 'Color', 'b'); %0.1X REF
hold on
plot(Xvec,cell2mat(TH{29,2}),Xvec,cell2mat(TH{30,2}), 'Color', 'b'); %0.01X REF
hold on


%adjust plot properties
ylim([0 6]);
xlim([5 35])
xticks(0:5:40);
xlabel('Cycle');

%%

subplot(3,5,5) %This is the linear amplification data
Xvec=transpose(1:40);

%plot all of the reference curves
plot(Xvec,cell2mat(REF{41,1}),Xvec,cell2mat(REF{42,1}), 'Color', 'r'); %1X REF
hold on
plot(Xvec,cell2mat(REF{43,1}),Xvec,cell2mat(REF{44,1}), 'Color', 'r'); %0.1X REF
hold on
plot(Xvec,cell2mat(REF{45,1}),Xvec,cell2mat(REF{46,1}), 'Color', 'r'); %0.01X REF
hold on
plot(Xvec,cell2mat(REF{47,1}),Xvec,cell2mat(REF{48,1}), 'Color', 'r'); %0.001X REF
hold on
plot(Xvec,cell2mat(REF{49,1}),Xvec,cell2mat(REF{50,1}), 'Color', 'r'); %0.001X REF
hold on

%plot all of the theta curves
plot(Xvec,cell2mat(TH{25,1}),Xvec,cell2mat(TH{26,1}), 'Color', 'b'); %1X REF
hold on
plot(Xvec,cell2mat(TH{27,1}),Xvec,cell2mat(TH{28,1}), 'Color', 'b'); %0.1X REF
hold on
plot(Xvec,cell2mat(TH{29,1}),Xvec,cell2mat(TH{30,1}), 'Color', 'b'); %0.01X REF
hold on

%adjust plot properties
xlim([10 40])
xticks(10:5:40);
xlabel('Cycle');
title('Time 5');


linkaxes([ax1 ax2 ax3 ax4 ax5],'xy')

toc

end
```

### 6. _Bacterial Biofilm imaging._

Wild type and PolYB- B. subtilis strains with a theta replicating plasmid were grown overnight at 37C. In the morning, cells were diluted 1:100 times and applied to agarose based microscope slides. After 12 hours of incubation at 37C in a sealed contained with 100% humidity, microscope slides were imaged using tritc filter set. Raw pixel intensity values were extracted from each image, and a histogram plot was created (Figure 8). Below is the code used to process these images:

```
function SporeImg
%Coded by NFR on 11.14.2018
% This code will help us process the spore images
% TB = Theta B
% 168 = wildtype
% PS prefix files = POST spore
%
% Concantanate all of the matrices initially
close all
TB = zeros(1040*5,1392*4);
c168 = zeros(1040*5,1392*4);
PS_TB = zeros(1040*5,1392*4);
PS_c168 = zeros(1040*5,1392*4);
for i = 1:20
    I1 = imread(['168_',int2str(i),'.tif']);
    I2 = imread(['TB_',int2str(i),'.tif']);
    I3 = imread(['PS_168_',int2str(i),'.tif']);
    I4 = imread(['PS_TB_',int2str(i),'.tif']);
    Id1 = double(I1);
    Id2 = double(I2);
    Id3 = double(I3);
    Id4 = double(I4);
    % Concantanate as a 5x4 composite image
    nr = ceil(i/4);
    nc = i-(nr-1)*4;
    TB(nr*1040-1039:nr*1040,nc*1392-1391:nc*1392) = Id1;
    c168(nr*1040-1039:nr*1040,nc*1392-1391:nc*1392) = Id2;
    PS_TB(nr*1040-1039:nr*1040,nc*1392-1391:nc*1392) = Id3;
    PS_c168(nr*1040-1039:nr*1040,nc*1392-1391:nc*1392) = Id4;
end

figure
histogram(PS_c168(:),'FaceColor', 'red', 'EdgeColor', 'red')
hold on
histogram(PS_TB(:))
xlabel('Pixel Intensity')
ylabel('Count')
xlim([0 3500])
ylim([0 3e5])
legend('168', 'PolYB-')
title('168 Before Sporulation')

figure
histogram(PS_c168(:))
xlabel('Pixel Intensity')
ylabel('Count')
xlim([0 3500])
ylim([0 3e5])
title('168 After Sporulation')

figure
histogram(TB(:))
xlabel('Pixel Intensity')
ylabel('Count')
xlim([0 3500])
ylim([0 3e5])
title('Theta-B Before Sporulation')

figure
histogram(PS_TB(:))
xlabel('Pixel Intensity')
ylabel('Count')
xlim([0 3500])
ylim([0 3e5])
title('Theta-B After Sporulation')
% Display scaled image with custom color map;
rvec = linspace(0,1,1000)';
zvec = zeros(1000,1);
map = [rvec zvec zvec];
figure
%imagesc(c168,[0 3500]) % This is 168 before spo montage
imagesc(c168,[0 3500])    % This is B- before spo montage
colormap(map);
figure
%imagesc(PS_c168,[0 3500]) % This is 168 after spo montage
imagesc(PS_c168,[0 3500])    % This is B- after spo montage
colormap(map);

%picking images to zoom in on
%B before is image 6
 bBefore = imread(['TB_',int2str(1),'.tif']);
 bBefore = bBefore(400:699, 400:699);
 imagesc(bBefore,[0 2000]);
%B after is image 6.
 bAfter = imread(['PS_TB_',int2str(2),'.tif']);
 bAfter = bAfter(400:699, 400:699);
 imagesc(bAfter,[0 2000]);
%168 before is image 11
before168 = imread(['168_',int2str(2),'.tif']);
before168 = before168(400:699, 400:699);
 imagesc(before168,[0 2000]);
%168 after is image 1
 after168 = imread(['PS_168_',int2str(1),'.tif']);
 after168 = after168(400:699, 400:699);
 imagesc(after168,[0 2000]);
end
```

### 7. Riboswitch Design and structure predictions.

Riboswitches were designed using the riboswitch calculator (attached <Riboswitch Parameters.xlsx>). Six riboswitches were designed, and three of them (Theo-43, -44, -45) were selected for testing (reference the xcel file). Riboswitch Theo-45 showed the highest level of activation (Fig. 1D), and was used for all of the growth experiments. The structure of the riboswitch Theo-45 was determined using MFold software (Supplement Figure 13).

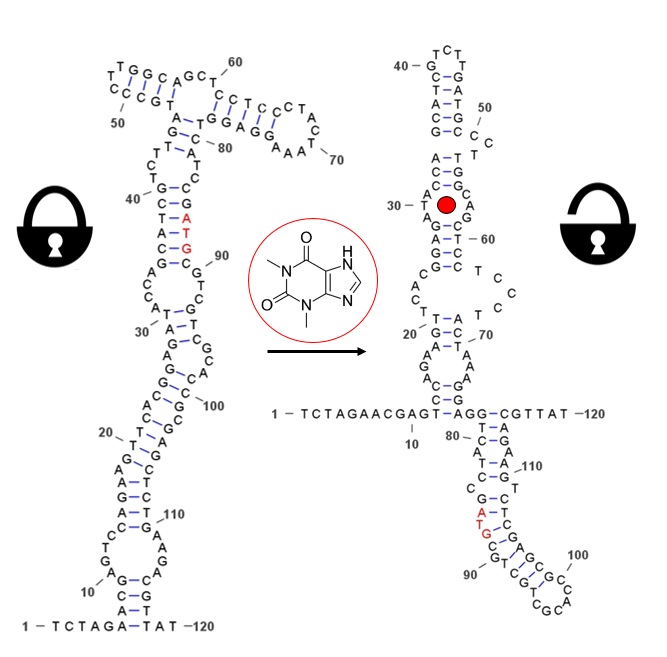

Supplement Figure 12 2D structure of the Theo-45 riboswitch generated using MFold.
